# Bioengineered bone marrows that can support stem cells and remodel to mimic human disease metabolism and chemotherapeutic effects

**DOI:** 10.64898/2025.12.03.692012

**Authors:** Yinbo Xiao, Savvas Ioannou, Monica P Tsimbouri, Xiang Li, Oana Dobre, Sara Trujillo, Mark Sprott, Mariana AG Oliva, Vineetha Jayawarna, Massimo Vassalli, Ewan Ross, Mhairi Copland, Peter S Young, RM Dominic Meek, Manuel Salmeron-Sanchez, Hannah Donnelly, Matthew J Dalby

## Abstract

Blood cancer drug discovery is reliant on poorly predictive and costly animal models. Therefore, bioengineered, human cell containing, models are attractive to Pharma. Designing effective bone marrow (BM) models is complicated as they need to both regulate haematopoietic stem cell (HSC) phenotype and be able to undergo remodelling to mimic the blood cancer microenvironment. Here, we develop synthetic hybrid niches using poly(ethylacrylate) to organise laminin on hard, bone mimicking, surfaces and interface with soft, marrow mimicking, polyethylene glycol-fibronectin hydrogels. Optimisation of mesenchymal stromal cell (MSC) mechanobiology within the model offers both support for HSCs and remodelling in response to acute myeloid leukaemia derived cells. The remodelling has many parallels to *in vivo* and patient data including increased dependency on nestin^+ve^ MSCs, enhanced cytoprotection and increased taurine metabolism. We also use the model to demonstrate, as has been seen in vivo, that targeting taurine enhances effects of chemotherapy.

---

There is a push for new chemotherapeutics for blood cancers. This push is slowed because animal models are slow (months to years), expensive, lack human specificity and can’t be used for high throughput screening. Human cell containing bioengineered models of blood cancer, therefore, have advantages of human physiology, speed (days to weeks), cost and throughput; if good models existed.

Blood cancers evolve in the bone marrow (BM) niche, a specialised local microenvironment in which blood-forming haematopoietic stem cells (HSCs) primarily reside^1^. BM composition is complex, dynamic and can remodel with malignancy to support diseased cancer stem cells over healthy HSCs^2,3^. This complexity presents barriers to the bioengineering of *in vitro* BMs. The BM contains two putative stem cell niches, the endosteal/arteriola (bone-lining) niche, responsible for LT-HSC maintenance and quiescence, and the sinusoidal, perivascular, niche, which is located in the central marrow cavity where HSCs are considered to divide and differentiate. Niche composition is highly heterogeneous; ECM consists mainly of collagen, fibronectin (FN), vitronectin and laminin (LM)^4,5^. FN is present throughout the BM, while collagen I is associated with the bone matrix, LM is distributed along the vascular basement membranes of arterioles and sinusoids^5^. ECM proteins are thought to have a strong influence on HSCs via stromal cells, such as mesenchymal stromal cells (MSCs). MSCs attach to the ECM and are influenced by ECM connecting integrin transmembrane receptors to release paracrine signals (e.g. MSC-derived CXCL12 (C-X-C motif chemokine 12), SCF (stem cell factor)) and to provide cell-cell adhesion molecules to support the HSCs^6–8^. The importance of the ECM can be seen from the effects of LM depletion *in vivo*, which induces engraftment failure post HSC transplantation^9^. MSCs are also influenced by mechanical properties of the BM. The BM is a heterogeneous hydrogel, it’s Young’s modulus varies from 0.3 to 24.7 kPa^10^; hydrogels with a Young’s modulus of ∼1-2 kPa induce MSCs to express the niche marker, nestin^6,11–13^.

Studies are emerging that offer insight. Bioengineered models based on ceramics to mimic the bone trabeculae and on on-chip systems have focused on murine HSCs or expansion of HSCs generally (including LT-HSCs but also their progeny - short term (ST)-HSCs and progenitor cells)^14–23^. A more recent study used collagen hydrogels overlayed on MSCs cultured on a FN interface and showed maintenance of human LT-HSCs^24^. While these studies help show us the way to maintain HSCs *ex vivo*, the second key function of a minimal model is missing, ability to remodel with disease development alongside data verified to *in vivo* and human disease.

Here, we optimise a synthetic-ECM interface, to resemble the endosteal surfaces, with hybrid synthetic-biological hydrogels, to represent the soft marrow. We build BM models that are tuned to support LT-HSCs. Then, for the first time, we show that our engineered niches can remodel to mimic the metabolic and cytoprotective effects of the leukemic niche that is observed in *in vivo* models and in human patients, and can accurately recreate drug responses seen *in vivo* that are missed in simple culture systems.

## Materials used to organise the ECM at the synthetic endosteal surface

A first aim is to build a healthy niche in which to initiate cancer progression. Our model comprises a hard-surface / soft gel interface to mimic an endosteum-to-BM interface, in which MSCs grow on the endosteum to support healthy HSC regulation **(Fig. S1)**. We have previously demonstrated the use of poly(ethylacrylate) (PEA) to organise FN^25^. FN contains cryptic binding sites that are made available to cells through protein unfolding or applied tension, usually cell-driven process^26^. When FN is adsorbed onto PEA, it spontaneously extends, forming networks that reveal its cell adhesion, RGD (arginine, glycine, aspartic acid) motif, and heparin-binding domain, which can bind many families of GFs, presenting them to cells in synergy with integrin-associated signalling^25^. This system has been used to drive *in vitro* and *in vivo* bone healing through a solid-phase presentation of ultra-low doses of BMP2^25^. To allow for future applications of high-throughput screens, we use standard 96-well plates and plasma polymerise PEA as a nanoscale coating onto the plates. Here, we use FN and LM as MSC interfaces as they are niche associated, and have different proposed mechanotransductive properties^27^.

Wells were successfully PEA coated, confirmed by X-ray photoelectron spectroscopy (XPS) (**Fig. S2**). To study **FN** and **LM** adsorption, 20 µg/ml of FN, LM or a (50/50) blend was adsorbed onto the **pPEA** surface, and atomic force microscopy (AFM) was used to assess network formation.

As expected for FN, which forms dense networks^28,29^, the FN only coating resulted in dense networks and LM only adsorption did not result in networks (**Fig. 1A**). However, when FN was blended with LM, networks were again observed (**Fig. 1A**), suggesting LM incorporation within the fibrils. This incorporation was confirmed using gold-labelled antibodies against LM. The anti-LM antibody was absent on the FN only surface, present on the LM only control surface, and present on the FN/LM blended surface, showing that LM had formed part of the nanonetwork (**Fig. 1B**).

**Fig. 1.**
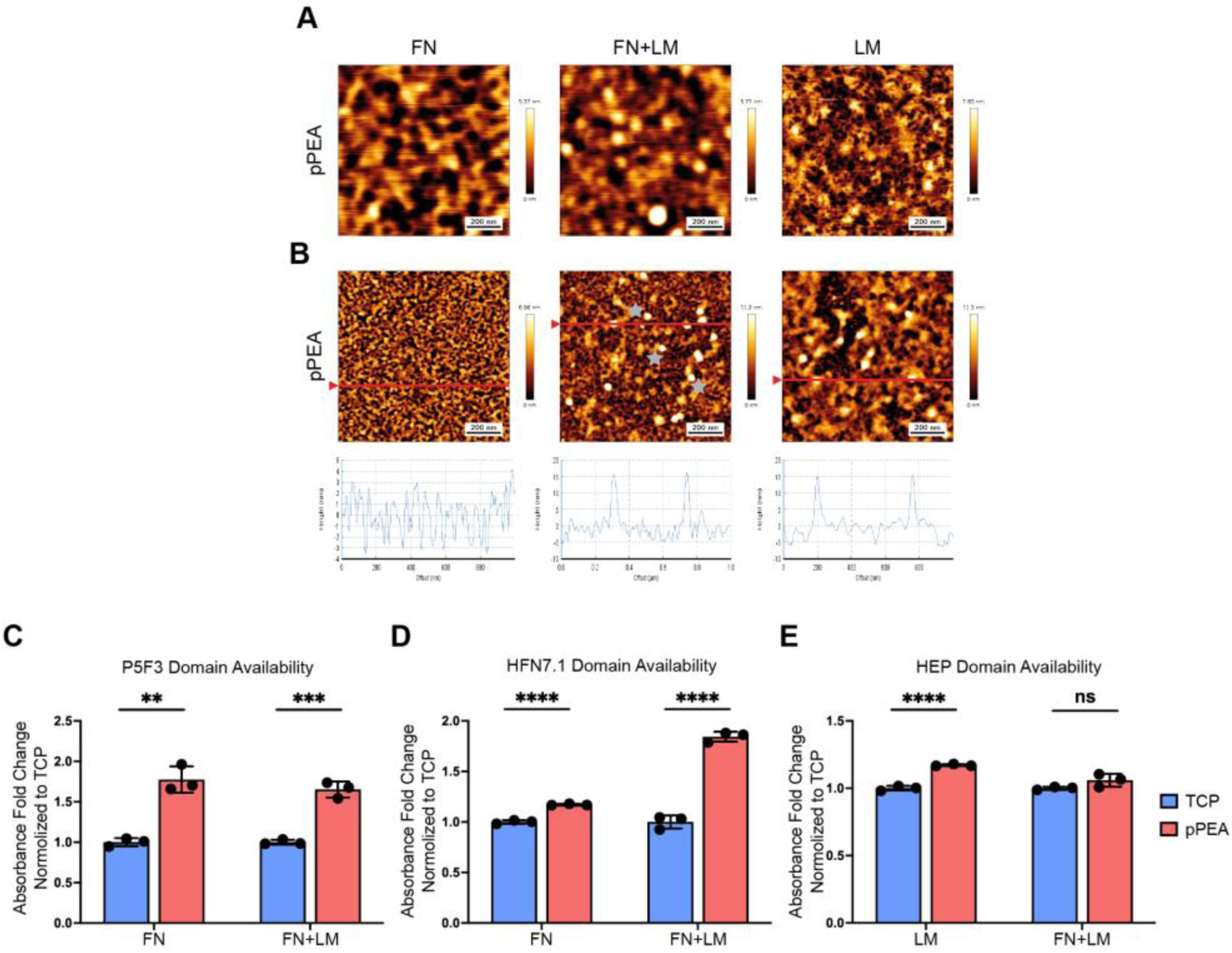
pPEA drives ECM network formation and functional domain exposure. **A,** AFM images of pPEA substrates coated with FN (left), FN+LM (middle), and LM (right). Scale bar: 200 nm. **B,** AFM images of LM incorporated into FN fibrillar nanonetworks on pPEA, where a secondary antibody bound to a 15 nm gold nanoparticle was used to identify LM. The measured height corresponding to image cross-sections (red line) is shown underneath each image to demonstrate antibody-nanoparticle locations. The LM incorporated within FN network is labelled with grey stars. Scale bar: 200 nm. **C, D, & E,** Availability of FN P5F3 domain (GF-binding domain) (**C**), FN HFN7.1 domain (RGD integrin-binding domain) (**D**), and LM HEP domain (GF-binding domain) (**E**) on TCP and pPEA surfaces. Mean absorbance fold change values over the TCP group with the same ECM coating were reported. Mean ± s.d. based on n= 3 material replicates. **, p<0.01, ***, p<0.001, ****, p<0.0001, ns, non-significant. P values were determined using the unpaired Student’s t-test.

Measurements of the amount of protein adsorbed showed that when FN was added to TCP or pPEA surfaces, significantly more FN adsorbed onto pPEA than TCP (**Fig. S3A&B**). When LM was added to the TCP or pPEA surfaces significantly more LM adsorbed on to pPEA compared to TCP (**Fig. S3A&B**). When a 50/50 FN/LM blend was added to pPEA or TCP, approximately half the amount of FN and LM were adsorbed on to pPEA and TCP compared to when they were added alone and there was no difference in absorption between pPEA and TCP (**Fig. S3A&B**).

To verify the availability of key functional sites, we quantified the exposure of three sites using specific antibodies. Both the cryptic FNIII_12–14_heparin-binding domain (anti-P5F3) and the RGD integrin binding site at FNIII_9-10_ (anti-HFN7.1) were significantly more available on the pPEA surface compared to the TCP surface for both FN and FN/laminin (LM) coatings (**Fig. 1C&D**). The heparin/GF binding site on LM (anti-HEP) was more available on pPEA than TCP (**Fig. 1E**). In the FN/LM blend, FN was seen to out-compete LM (**Fig. 1C-E**). These results indicate that FN adsorbs onto 2D pPEA-coated surfaces in an open network configuration with available key functional domains, allowing LM to integrate. Although LM alone doesn’t form a fibrillar network on pPEA, its amount and GF binding domain availability are increased on pPEA relative to TCP.

## Optimising mechanotransductive and GF interactions at the endosteal interface to achieve stromal phenotype

We next investigated differences in human MSC adhesion and spreading in response to ECM surfaces. After 3 hours culture, actin cytoskeleton and vinculin located at adhesions were stained, and projections of cell adhesions were processed (**Fig. 2A**, **Fig. S4A**) and cell area calculated (**Fig. 2B**). There was no difference in cell area for pPEA coated with FN, FN/LM or LM (**Fig. 2B**). However, cells spread more on pPEA-coated surfaces, than on TCP (**Fig. S4B**). On pPEA, the addition of LM alone or blended with FN, reduced both adhesion number (**Fig. 2C**) and cell spreading (**Fig. 2D**). LM also decreased the numbers of larger adhesions in favour of smaller adhesions (**Fig. 2E**, **Fig. S4C&D**).

**Fig. 2.**
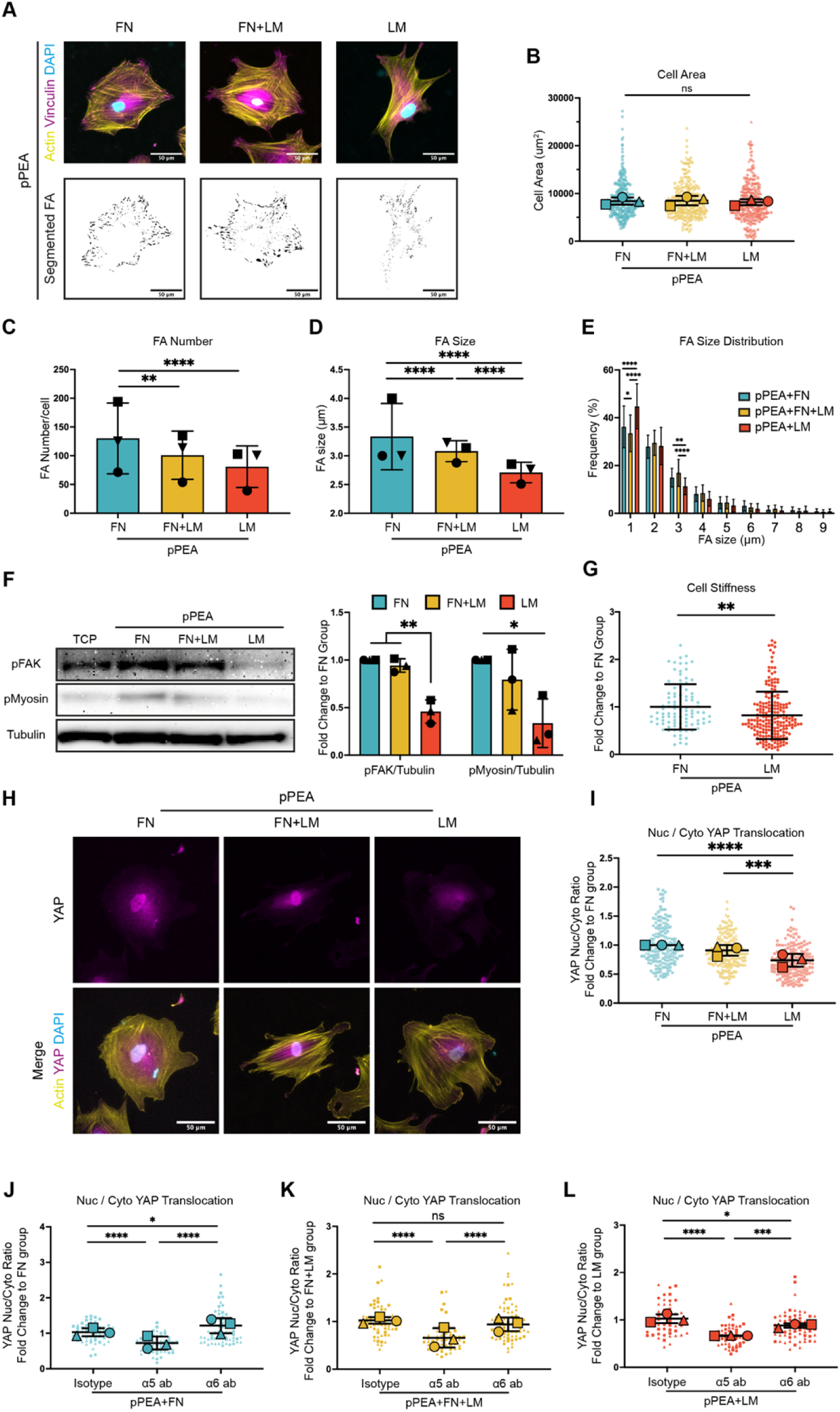
LM reduces MSC mechanotransduction. **A,** Representative immunofluorescence images of vinculin(magenta)/ actin(yellow)/ DAPI(blue) stained, and thresholded binary images for MSCs’ FA analysis. Scale bar: 50 μm. **B, C, D, & E,** analysis of cell area **(B)**, focal adhesion (FA) number **(C)**, FA size **(D)**, and FA size distribution **(E)** based on actin and vinculin staining. **F,** Western blot analysis of the phosphorylation level of FAK and myosin in MSCs cultured on FN, FN+LM, and LM pPEA surfaces. **G,** Stiffness (Young’s modulus by nanoindentation) of MSCs cultured on FN and LM pPEA surfaces. **H,** Representative immunofluorescence images of YAP (magenta)/ actin (yellow)/ DAPI (blue) staining of MSCs on FN, FN+LM and LM pPEA surfaces. Scale bar: 50 μm. **I,** Quantification of YAP nuclear/cytoplasmic (nuc/cyto) ratios in MSCs culture on FN, FN+LM LM pPEA surfaces. **J, K, & L,** Quantification of YAP nuc/cyto ratios in MSCs cultured on FN **(J)**, FN+LM **(K)**, and LM **(L)** pPEA surfaces after blocking α5 or α6 integrin with neutralising antibodies. Each shape represents each donor, each small shape represented each cell, and each large shape represents the mean from each donor. Mean ± s.d. based on n= 3 independent experiments with different donor cells. *, p<0.05, **, p<0.01, ***, p<0.001, ****, p<0.0001, ns, non-significant. P values were determined by unpaired Student’s t-test (**G**) and ordinary two-way ANOVA with Tukey’s multiple comparisons test (**B-F, I-L**).

Activation (phosphorylation) of the mechanotransductive signalling protein focal adhesion kinase (FAK)^30,31^ in cells on FN and LM adsorbed to pPEA was assessed. We observed similar levels of FAK activation in MSCs on all surfaces except pPEA-adsorbed LM, where it was reduced (**Fig. 2F and Fig. S5**). This suggests that FN might be dominant to LM in terms of mechanotransduction. We next investigated myosin activation (phosphorylation), cell stiffness and YAP (yes associated protein) nuclear translocation as markers of mechanotransductive changes^32,33^, with levels of intracellular tension lower than on TCP linked to retention of the MSC phenotype^34–36^. We observed reduced levels of mysosin activation (**Fig. 2F**), cell stiffness (**Fig. 2G**) and YAP translocation to the nucleus (**Fig. 2H,I**) in cells on pPEA-adsorbed LM compared to those on pPEA-adsorbed FN. This all suggests that LM reduces the mechanotransducive state of the MSCs compared to FN, but that FN is a dominant matrix cue.

Our data fits with literature on epithelial cells showing that LM reduces intracellular tension, lowering YAP translocation, while maintaining adhesion^27^. To test this further, MSCs were exposed to blocking antibodies for integrins a_5_ (FN receptor) or a_6_ (LM receptor) and YAP translocation was measured. For MSCs on pPEA-adsorbed FN, FN/LM, or LM, blocking the FN receptor decreased YAP nuclear translocation relative to isotype control, showing that FN is important for sustaining mechanotransduction (**Fig. 2J-L**). By contrast, blocking the LM receptor increased YAP translocation, indicating that LM attenuates mechanotransduction (**Fig. 2J-L**, **Fig. S6**).

This lowered mechanotransductive state should support a more stromal MSC phenotype^36^. Stromal MSCs express high levels of nestin and are more secretory. MSCs on pPEA-adsorbed LM retained their expression of nestin (**Fig. 3A**) and released a range of GFs that are involved in HSC support^8^, including CXCL12, osteopontin (OPN), vascular cell adhesion molecule (VCAM) and vascular endothelial GF (VEGF). These MSCs also released several immunomodulatory factors, including interleukin 6 (IL-6), MIF (macrophage migration inhibitory factor) and IGFBP3 (insulin-like GF binding protein 3)^37^ (**Fig. 3B & Fig. S7**). Principal component analysis confirmed that the cytokine/GF profile of MSCs on LM, FN, and FN/LM are distinct (**Fig. 3C**). Overall, LM maintains a stromal MSC phenotype, marked by nestin and specific cytokine/GF expression, by modulating MSC mechanotransduction and, therefore, was selected as our ECM interface.

**Fig. 3.**
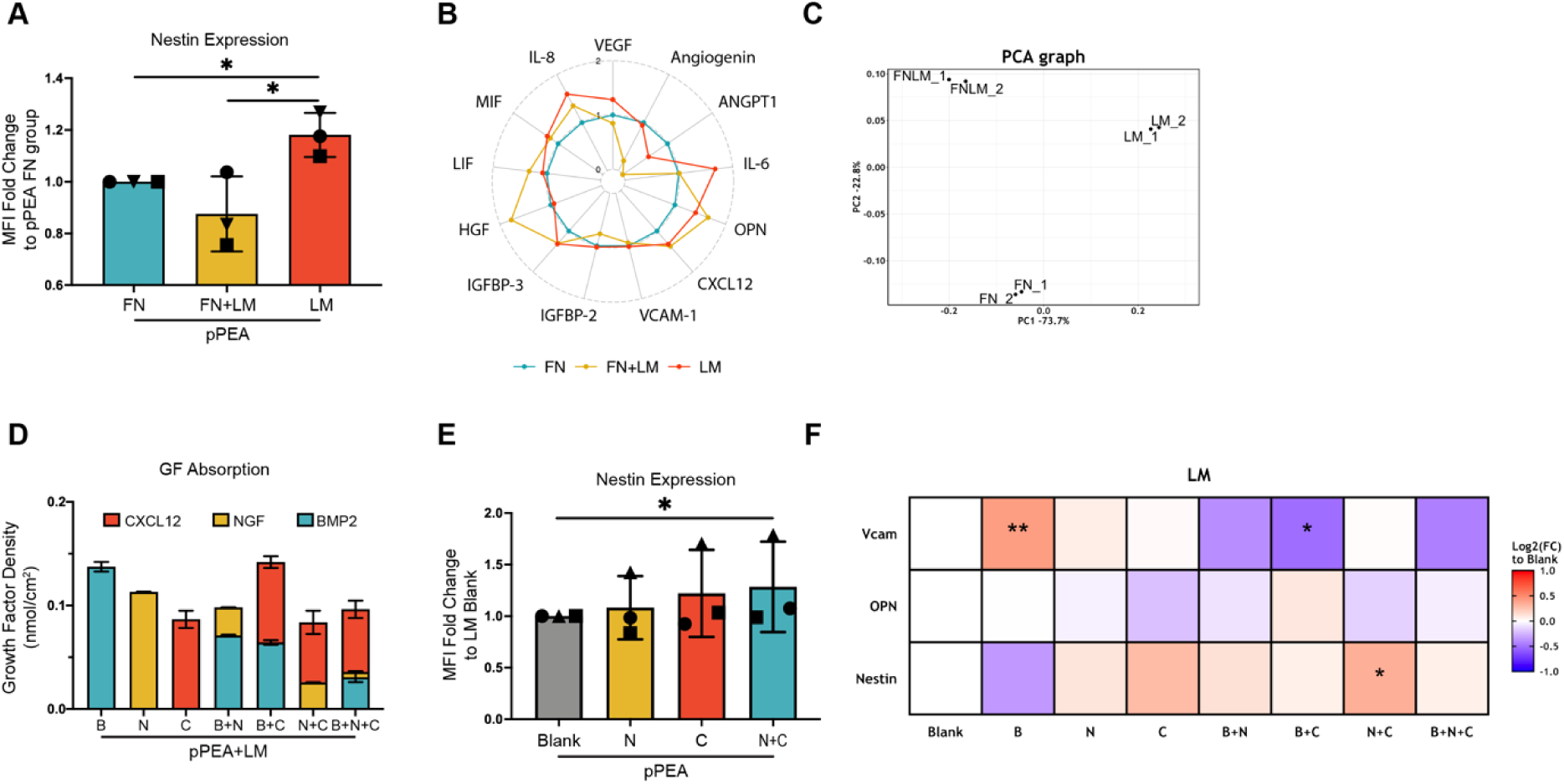
LM maintains MSC niche phenotype and LM+NGF+CXCL12 further enhance phenotype. **A,** In-cell western analysis of nestin expression of MSCs cultured on FN/FN+LM/LM pPEA surfaces for 14 days. Each shape represents each donor. **B,** Radar plot representing the GF secretion profile of MSCs cultured on FN/FN+LM/LM pPEA surfaces for 14 days. Data shows the MFI fold change values over the FN group (colours represent the same groups as used in panel A). **C,** PCA analysis of GF secretion profile of MSCs cultured on FN, FN/LM and LM pPEA surfaces. **D,** ELISA analysis of GF absorption density on pPEA/LM surface. **E,** nestin expression of MSCs cultured on blank, NGF, CXCL12, and NGF/CXCL12 pPEA-LM surfaces. Each shape represents each donor. **F,** Heatmap representing nestin, OPN and VCAM expression for MSCs cultured on the pPEA/LM/GF surface for 14 days. In-cell western analysis was utilised. Log_2_-MFI fold change values over the pPEA/LM group was reported. **A, E&F**, Mean ± s.d. based on 3 independent experiments with different donor cells. **B&C**, Based on one independent donor with technical duplicates. **D,** Mean ± s.d. of n= 3 material replicates. *, p< 0.05, **, p< 0.01. ns, non-significant. P values were determined by ordinary two-way ANOVA (**A, E&F**). (B: BMP2; N: NGF; C: CXCL12; B+N: BMP2/NGF; B+C: BMP2/CXCL12; N+C: NGF/CXCL12; B+N+C: BMP2/NGF/CXCL12).

Next, to further optimise our endosteal mimic, we assayed for the binding of a range of GFs to the cryptic binding sites that are exposed in pPEA-adsorbed LM, and specifically for BMP2, CXCL12 and neural GF (NGF) as possible niche factors due to their presence at endosteal bone surfaces^38^. ELISA confirmed that all tested GFs, individually and combined, bind to pPEA-adsorbed LM and remain available to cells (**Fig. 3D**). In-cell western analysis of MSC expression of niche factors when grown on the GF loaded surfaces (nestin, VCAM, OPN) showed that BMP2 generally lowered nestin (alone) and VCAM (in combination with other GFs) expression and so was dropped (**Fig. 3E**). CXCL12 and NGF appeared to up-regulate VCAM and significantly up-regulated nestin (**Fig. 3E**). The highest nestin expression was achieved with the combined CXCL12 and NGF (**Fig. 3F**). Therefore, the pPEA/LM/NGF-CXCL12 (NC) combination was selected as the optimised niche surface for subsequent experiments.

## Building the haematopoetic niche with synthetic-biological hybrid hydrogels

The BM is an ECM-rich hydrogel, and we know that soft hydrogels cause high nestin expression levels in MSCs^13^. We want reproducible, synthetic, hydrogels that also confer ECM functionality by, for example, acting as a reservoir for cytokines and GFs. We therefore developed polyethylene glycol (PEG) hydrogels in which stiffness is controlled via the ratio of PEG and crosslinkers. We incorporate matrix metalloprotease (MMP) cleavable VPM (GCRD**<u>VPM</u>**SMRGGDRCG) peptide crosslinkers to allow recovery of cells for analysis. The hydrogels are 3% PEG-maleimide (PEG-MAL) crosslinked with (PEG-dithol (PEG-SH) or VPM (75:25% VPM:PEG-SH) crosslinker ratio and a BM-like stiffness of ∼3 kPa (**Fig. 4A**) ^39^. To give biological functionality, we added PEGylated FN to the hydrogel network to allow recapitulation of the GF capture properties of natural hydrogels^39^. The addition of PEGylated FN reduced gel stiffness to ∼2 kPa (**Fig. 4A**) and increased the gels ability to adsorb and retain niche-related GFs, such as CXCL12 (**Fig. 4B&C**) and SCF (**Fig. S8A&C**). Some GFs, such as FMA-like tyrosine kinase 3 ligand (Flt3L), were less well adsorbed (**Fig. S8B&D**).

**Fig. 4.**
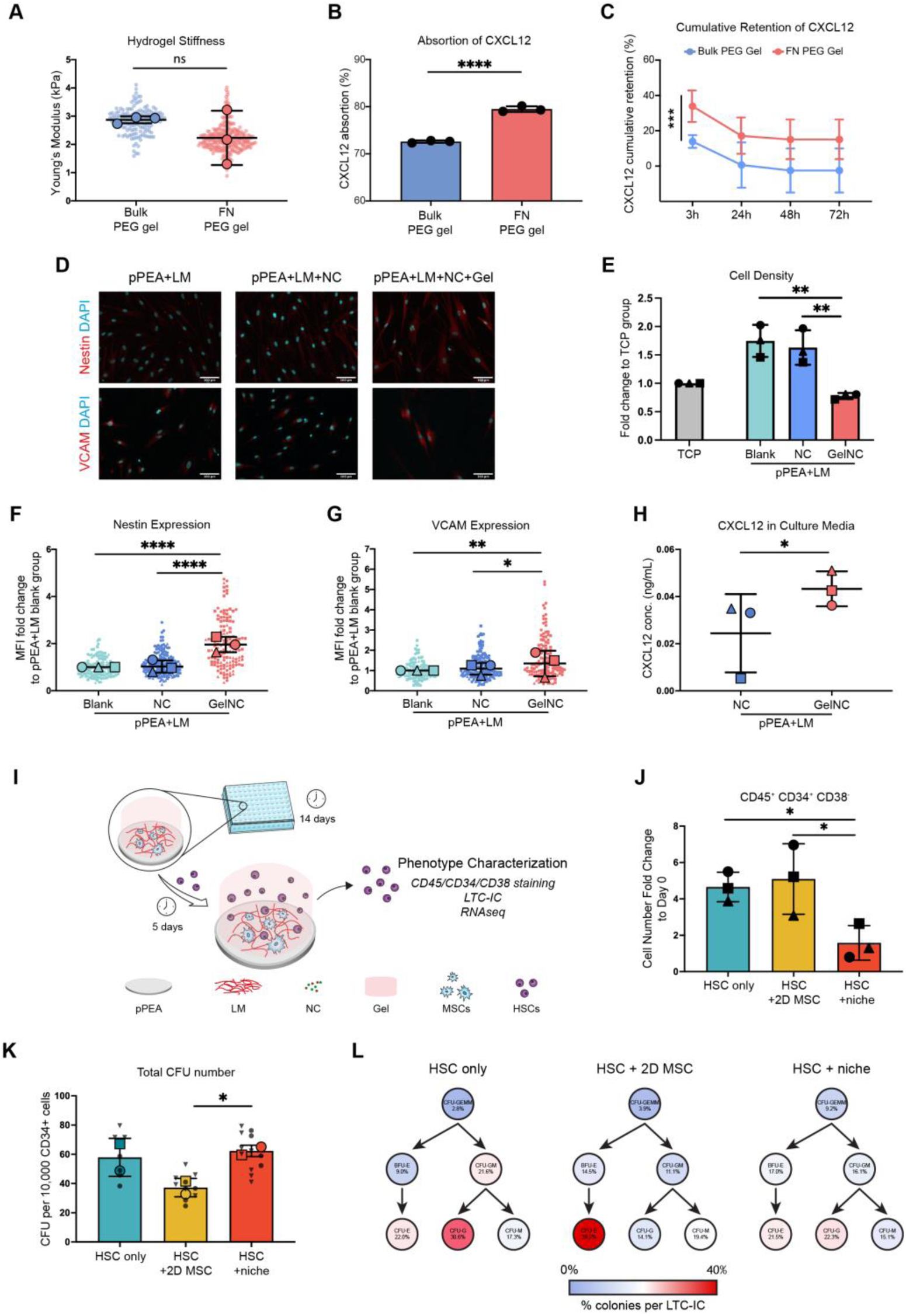
Hydrogels help mimic the bone marrow. **A,** The Young’s modulus of bulk PEG hydrogels and FN PEG hydrogels. At least 100 curves were obtained from each hydrogel. **B,** The percentage of adsorbed CXCL12 in the bulk PEG gel and the FN PEG gel. **C,** The percentage of cumulative CXCL12 retained in the bulk PEG gel and the FN PEG gel. **D,** Representative images of nestin and VCAM staining in MSCs within the niche model. Scale bar: 100μm. **E,** MSC number within the niche models. **F&G,** Immunofluorescence of Nestin and VCAM expression of MSCs cultured in the FN-PEG gels with NC (GelNC) /without (NC) for 14 days. Quantification analysis of MFI fold change values over the pPEA+LM (Blank) group is presented. **H,** ELISA analysis of CXCL12 production by MSCs within the niche models. (**I**) Schematic of HSC co-culture in the niche model. **J,** Cell count for HSCs (CD45^+^CD34^+^CD38^-^) cultured in the niche models for 5 days. The cell number fold change to the day 0 input is presented. **K,** Total number of colonies generated in LTC-IC assays from each niche model. **L,** Lineage differentiation of LTC-ICs, as demonstrated by colony type, from the niche models. The colour charts are based on the percentage (%) of CFU-GEMM, BFU-E, CFU-GM, CFU-E, CFU-G, CFU-M of total CFU colonies counted in each model and arranged by haematopoietic hierarchy. Each shape represents individual donors, each small shape represents individual cells, and each large shape represents the mean from each donor. **A-C,** Mean ± s.d. based on n= 3 material replicates. **E-H,** Mean ± s.d. based on n= 3 independent experiments with different donor cells. *, p<0.05, **, p<0.01, ***, p<0.001, ****, p<0.0001, ns, non-significant. P values were determined by unpaired Student’s t-test (**A, C, H**), paired Student’s t-test (**B**) and ordinary two-way ANOVA with Tukey’s multiple comparisons test (others). (NC: NGF+CXCL12).

First, we assessed MSC number, nestin and VCAM expression at the hard-surface / soft-gel interface in the NC (NGF and CXCL12) system, without and with FN-PEG gel (GelNC), using image analysis at day 14. The addition of the GelNC slowed MSC growth (**Fig. 4D&E**) and increased expression of nestin and VCAM (**Fig. 4F&G**). ELISA showed increased CXCL12 expression at day 14 (**Fig. 4H**). OPN expression was reduced with addition of the gel (**Fig. S8E&F**). Therefore, our stromal niche comprises pPEA adsorbed LM, decorated with NGF and CXCL12, and overlayed with a degradable FN-PEG hydrogel with MSCs at the pPEA-hydrogel interface.

Next, we investigated HSC growth and phenotype when added to MSC-preconditioned synthetic niches. MSCs were cultured for 14 days using DMEM with 2% FBS before adding primary human CD34+ HSC/progenitor cells. The medium was then switched to gold-standard HSC media (IMDM with Flt3, SCF, and thrombopoietin, TPO) (**Fig. 4I**). This step is necessary to demonstrate niche functionality by supporting the long-term HSC (LT-HSC) population.

We used flow cytometry to assess the growth and phenotype of HSCs. We gated for CD45^+^CD34^+^CD38^-^ cells (LT-HSCs, ST-HSCs (stem) and primitive progenitor cells, together termed haematopoietic stem and progenitor cells (HSPCs) recovered from hydrogels post-culture (**Fig. S9A**) and observed that this population had expanded in the gold-standard media controls and in the 2D MSC culture systems (pPEA/LM/NC) (**Fig. 4J**). However, in the 3D synthetic niche, this population’s cell number remained the same as at day 0 (**Fig. 4J**). CD45^+^CD34^+^CD38^+^ (haematopoietic progenitor cells, HSPCs) and CD45^+^CD34^-^CD38^+^ (committed progenitor) populations did not expand in any condition (**Fig. S9B&C**), illustrating that any expansion remains in the primitive CD45^+^CD34^+^CD38^-^ HSPC population.

We next used the long-term culture initiating cell assay (LTC-IC) to identify true LT-HSCs. Lin^-^CD34^+^ cells were recovered from the controls and niche (**Fig. S10**) and placed into defined culture conditions for over 6 weeks followed by colony-forming unit (CFU) assay. This enables assessment of LT-HSCs colony forming potential. Our data shows that the synthetic niche maintained the largest population of LT-HSCs, compared to HSCs in gold-standard media and in 2D MSC co-culture (**Fig. 4K**). Together with our flow cytometry data (**Fig. 4J**), this illustrates that HSPCs are expanding in the gold-standard condition at the expense of LT-HSCs. Phenotyping of differentiated LT-HSC colonies revealed that while LT-HSCs cultured in gold-standard media and over 2D MSCs exhibited lineage bias towards granulocytes and erythrocytes, respectively, LT-HSCs cultured in the synthetic niche exhibited less lineage bias (**Fig. 4L & Fig. S11**). Together, these data show that LT-HSCs in the synthetic niche are maintained without expansion of ST-HSCs and progenitors, and without developing unwanted lineage bias.

## MSC remodelling in the leukemic niche

To model leukaemia in our synthetic niche, we cultured MSCs for 14 days, and then added THP1 cells (an immortalised human monocytic AML cell line) for 3 days to allow for niche remodelling (**Fig. 5A**). Using a frontline AML chemotherapeutic, cytosine arabinoside (AraC)^40^, we identified IC_50_ for THP1 monoculture viability (**Fig. S12**). We assayed mitochondrial ROS (mtROS) to measure cell stress (**Fig. 5B**) and used propidium iodide (PI) to assay for cell death (**Fig. 5C & Fig. S13**). We then compared THP1 cells in monoculture and in the synthetic niche, observing lower cell stress and cell death when THP1 cells were treated with AraC in the niche. We also observed that MSC nestin expression increased when MSCs were co-cultured with THP1 cells (AML niche) (**Fig. 5D&E**), and that MSC numbers increased (**Fig. S14**) compared to being cultured without THP1 cells (3D niche only); AML has been shown to reduce the viability of stromal cells that do not express nestin in BM *in vivo*^41^. It has also been proposed that AML cells use the nestin-expressing stroma to remodel the BM environment to the advantage of cancer cells^41,42^. To assess this, we stained human BM sections from healthy donors and AML patients and observed significantly more nestin+ cells in the BMs of AML patients (**Fig. 5F&G**). To investigate MSC activation in response to THP1 cells, we set up niche models with no HSCs (3D niche) or THP1 cells (AML niche), and also healthy niches containing normal human CD34^+^ cells, and then isolated the MSCs for RNA/metabolite extraction (**Fig. 5H**). Looking at whole-cell transcriptional profiles, we observed each condition was distinct (**Fig. S15**) and for MSCs from 3D niche, the addition of HSCs (healthy niche) caused a modest upregulation of gene transcription (**Fig. 5I**). However, the addition of THP1 cells caused large-scale changes in gene transcription (**Fig. 5I**). RNAseq linked KEGG analysis indicated that MSCs in the AML niche reduced ECM and cell-cell adhesion while increasing actin cytoskeleton regulation. Analysis also indicated reduction of senescence and increased growth, and up-regulation of pathways involved in MSC-HSC interactions, such as HIF and TGFb (**Fig. 5J**)^43,44^. Together, these findings suggest that the MSCs become less influenced by their matrix and more influenced by the cancer cells.

**Fig. 5.**
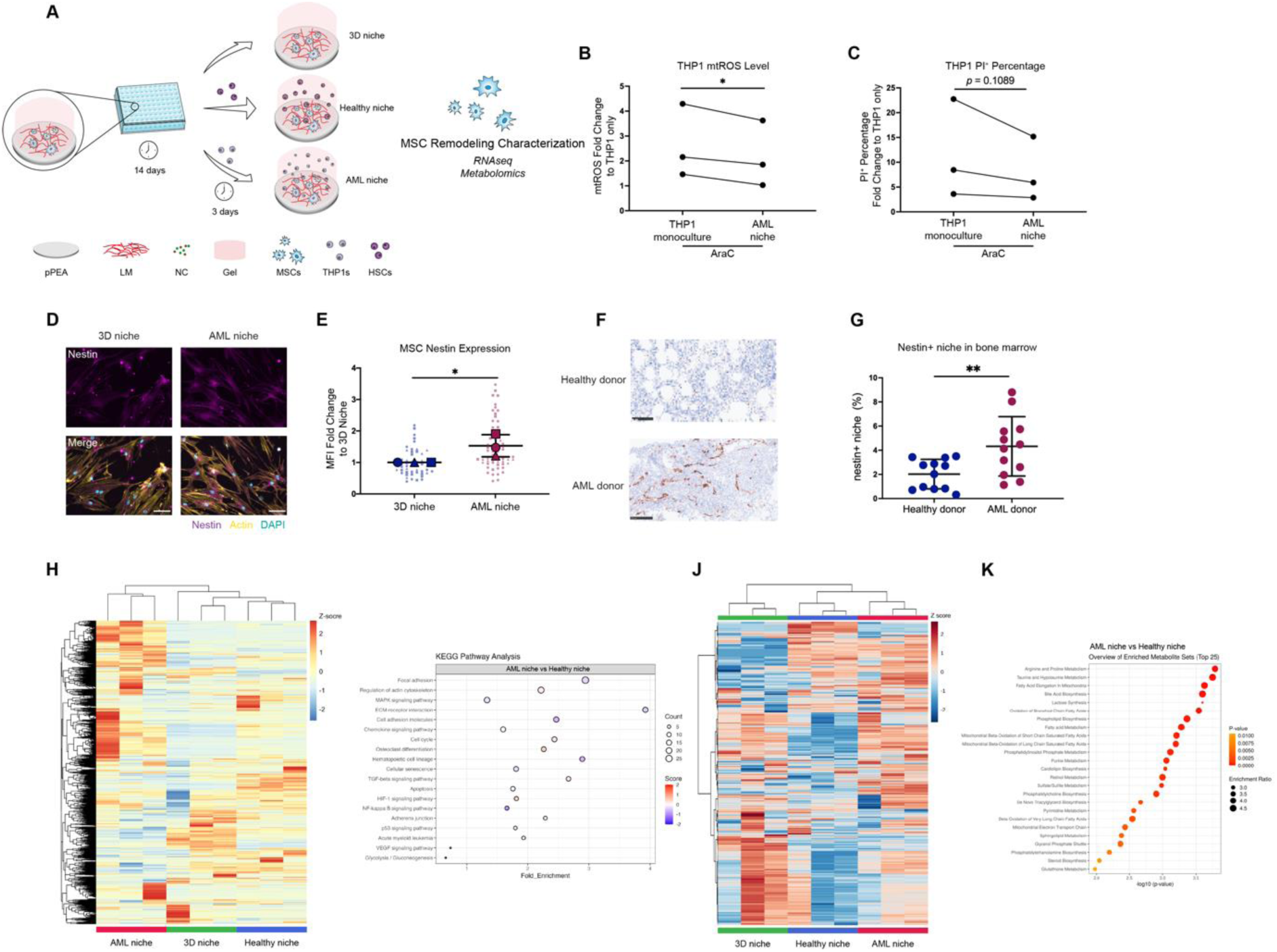
Cancer cells cause the niche to remodel. **A,** Schematic of leukemic cell co-culture in the niche model. **B,** mtROS analysis of THP1 from monoculture or in the niche model with AraC chemotherapy. **C,** PI analysis of THP1 from monoculture or in the niche model with AraC chemotherapy. Quantification analysis of fold change values over the THP1-only group without AraC treatment is presented. Graph shows mean from n= 3 independent experiments with different donor cells. **D,** Representative images of Actin (yellow)/ nestin (magenta)/ DAPI (blue) staining in MSCs from the 3D niche and AML niche models. Scale bar: 50 µm. **E,** Nestin expression of MSCs in the 3D and AML niche models. Quantification analysis of MFI fold change values over the 3D niche group are presented. **F&G**, Nestin^+^ area in bone marrow sections of healthy donors and AML donors (nestin is stained brown, scale bar is 100 µm). Nestin^+ve^ area as a percentage of 200x image field. Mean ± s.d. based on n= 12 different donors per group. **H,** Heatmap representing differentially expressed genes (DEGs) in MSCs from the different niche models. **I,** KEGG enrichment analysis of MSCs from the AML niche and the healthy niche. **J,** Heatmap of the metabolite changes in MSCs from the niche models. **K,** Enrichment analysis of metabolites with statistical differences in MSCs from AML niche and healthy niche. All graphs show 3 independent donors for each group. In superplots, each shape represents individual donor, each small shape represents individual cells, and each large shape represents the mean from each donor. In Fig. B, C and E, Mean ± s.d. based on n= 3 independent experiments with different donor cells. *, p<0.05; **, p<0.05. P values were determined by unpaired Student’s t-test (**E,G**), and paired Student’s t-test (**B, C**).

The metabolomes of MSCs in the healthy niche and in the AML niche exhibited many changes (**Fig. 5K & Fig. S16**). The pathways that show the most significant upregulation following the addition of THP1 cells included taurine, amino acid synthesis, respiration (mitochondria) and lipid pathway (**Fig. 5L, Fig. S17**). These trends fit well with the enhanced energy needs (lipids and respiration) and with the increased protein synthesis demands (amino acids) placed on activated MSCs. Taurine has also been implicated in an MSC ‘neural-like’ phenotype that features increased nestin expression^45^, and fatty acids have been implicated in supporting the survival and proliferation of AML blasts^46,47^.

## Using taurine metabolism to validate the bioengineered niche

Considering our data, that *in vivo* study has shown that taurine from the tumour microenvironment is implicated in AML progression^48^, and that taurine transporter loss-of-function mouse models have very recently shown that inhibition impairs *in vivo* myeloid leukemia progression^48^, we decided to focus on taurine in model validation. For all cell experiments, niches were primed for 14 days with MSCs before addition of HSCs/THP1s for 3 days. Using mass spectrometry from MSCs isolated from the healthy and AML niche models post addition of HSCs/THP1s, we looked at expression of taurine and taurine precursors L-cysteate, 3-sulfino-L-alanine and hypotaurine in the healthy or AML niche models; taurine, 3-sulfino-L-alanine and hypotaurine were all significantly elevated in the AML niche models (**Fig. 6A**). RNAseq data was mined for enzymes involved in taurine biosynthesis, including glutamate decarboxylases (GAD1,2) and GAD-like protein 1 (GADL1), cysteine sulfinic acid decarboxylase (CSAD), cysteine dioxygenase type 1 (CDO1), γ-glutamyltransferase 6 (GGT6) and flavin-containing monooxygenases (FMO1-5)^49–51^. Data showed increased expression of these enzymes in the AML niche model compared to the healthy niche model (**Fig. 6B**).

**Fig. 6.**
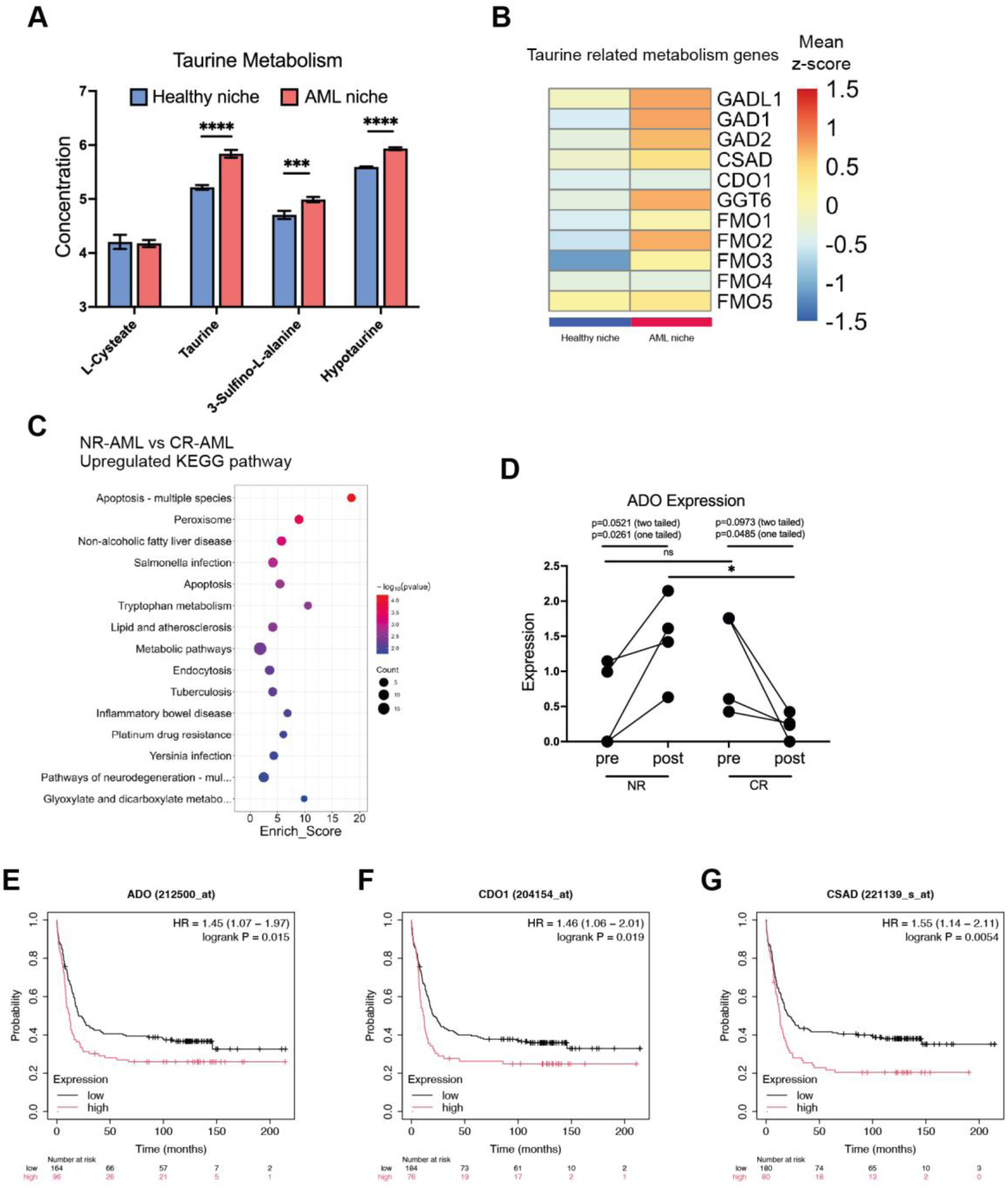
MSC taurine metabolism supports LSC chemoresistance in the clinic. **A**. Taurine-related metabolite changes in MSCs from the niche models. **B**. Taurine-related genetic changes in MSCs from the niche models. **C**, Proteomic analysis of bone marrow samples from NR and CR donors. Metabolic pathways were significantly different between NR and CR donors by KEGG enrichment analysis. **D**. ADO expression comparison between NR and CR donors. Based on n=4 donors per group. *, p<0.05; **, p<0.05, ns, non-significant. P values were determined by the paired Student’s t-test. **E-G**. Prognostic role of taurine metabolism-related genes in the AML GSE1159 database. Kaplan-Meier curve stratified by the expression of taurine metabolism-related genes. A log-rank test was used for comparisons.

Looking again at AML patient data, patients were grouped as non-responsive (NR) and complete-remission (CR) following AraC treatment. Proteomic analysis showed broad differences in metabolic pathways in the NR donors compared to the CR donors (**Fig. 6C**). Focusing on the expression of cysteamine dioxygenase (ADO), a key enzyme in taurine metabolism that converts cysteamine to hypotaurine ^52^, analysis showed that for NR patients, ADO levels significantly increased (**Fig. 6D**). This is suggestive of increasing taurine metabolism in response to chemotherapy. However, for CR patients, ADO levels reduced into remission (**Fig. 6D**). Employing the AML GSE1159 database (NIH AML gene expression database), it was seen that elevated expression of ADO, CDO1 (converts cysteine to cysteine sulfinic acid) and CSAD (converts cysteine sulfinic acid into hypotaurine)^49–51^, are linked to increased patient mortality (**Fig. 6 E-G**).

As discussed, a recent report on targeting taurine transporters *in vivo* showed inhibition of the TAUT (solute carrier family 6 member 6) taurine transporter inhibited AML progression^48^. Therefore, we used P4S (piperidinyl-4-sulfonic acid) as a competitive inhibitor of the taurine transporter to see effect in our AML niche model vs THP1 monoculture and THP1 with 2D MSC co-culture (P4S was added at the same time as the THP1s). We have the hypothesis, developed from *in vivo* data ^48^ and our human data (**Fig. 6 D-G**), that stromal cells secrete taurine and AML cells are supported by this to grow. In support of this hypothesis, we saw no effect of P4S addition on THP1 monoculture indicating stroma are required (**Fig. 7A**). With the no-gel, 2D (pPEA/LM/NC) co-culture, we saw a reduction in CFSE intensity that indicates increased THP1 growth; this is opposite to hypothesis (**Fig. 7B**). However, on hypothesis, in our 3D AML niche, we saw increased CFSE intensity linked to reduced THP1 growth (**Fig. 7C**). To test this further, we studied the additive effect of P4S with AraC. It was seen in monoculture, that there was no added benefit if P4S addition (**Fig. 7D**), With the 2D co-culture, we saw a highly significant increase in THP1 killing with use of P4S together with AraC (**Fig. 7E**). This enhanced killing remained in our 3D AML niche, but with reduced significance (**Fig. 7F**), showing that the inhibitor can enhance the efficacy of the front-line therapy, but that niche protection is still influential.

**Fig. 7.**
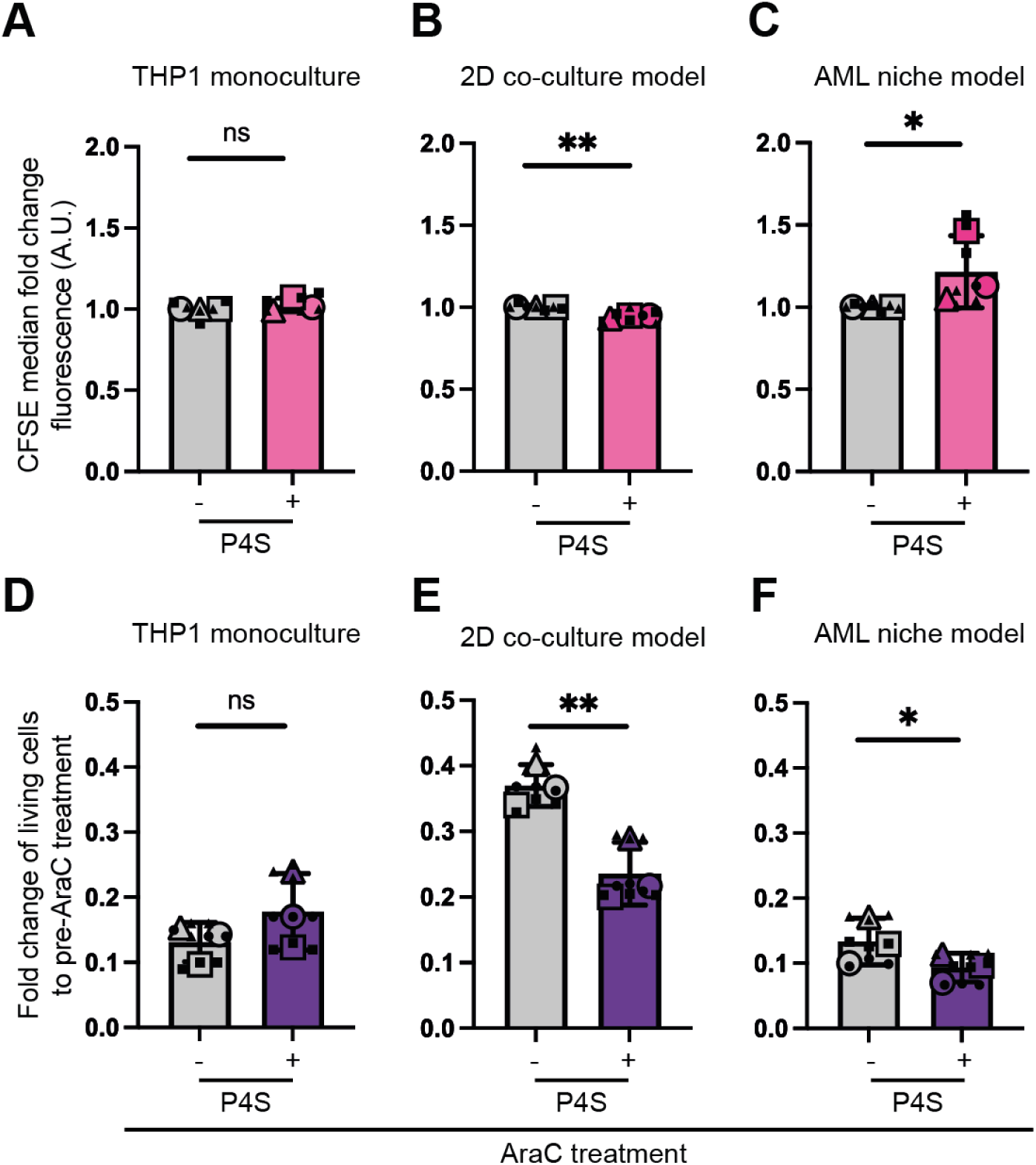
Testing anti-tumour efficacy of the Taurine inhibitor (P4S) in the AML niche model. **A-C.** AML cells were stained with CFSE, and their proliferation potential was assessed via flow cytometry after 72 hours in the presence or absence of P4S. **D-F.** AML cells were cultured under different conditions (monoculture, 2D co-culture, AML niche model) for 72 days, followed by AraC treatment for 72 hrs. Their viability was assessed by flow cytometry using the zombie violet stain. Each shape represents individual donors, each small shape represents technical replicates, and each large shape represents the mean from each donor. A-C P values were determined by lognormal Welch test. D-F P values were determined by paired Student’s t-test

Together, the data confirms that taurine, and related metabolism, is upregulated in our AML niche model, as it is *in vivo*^48^ and as it is in patients (**Fig. 6**). We then show that our model containing THP1 cells rather than HSCs, mimics data derived *in vivo* and highlights taurine as a therapeutic target. Monoculture of THP1 cells revealed very little in terms of investigating taurine and standard 2D co-culture gave either opposite or exaggerated results compared to our bioengineered BM model; both could give false hypothesis. We finally show that P4S can be used to augment the killing of AraC; this agrees with *in vivo* data showing inhibition of TAUT can increase efficacy of chemotherapeutics.

## Summary

The BM niche plays a dual role in health and disease, particularly in leukaemia where cancer cells remodel the microenvironment. Effective drug screening requires humanised (human cell containing) models that support LT-HSC maintenance and niche remodelling. Here, we present a synthetic-natural hybrid niche that meets these criteria. Using pPEA to organise LM, we modulate MSC mechanotransduction, enhancing nestin expression and secretion of HSC-supportive factors. MSC phenotype is further enhanced upon addition of a soft PEG hydrogel that contains FN as a GF sink, leading to enhanced support of human LT-HSCs.

For disease remodelling, we have a number of points of strong correlation from our bioengineered BM to *in vivo* and in human data. Firstly, using nestin as a stromal marker where *in vivo* study has shown nestin expression increases with SML disease progression;^41^ we reproduce this both in our model (**Fig. 5 D&E**) and in human samples (**Fig. 5 F&G**). Secondly, our bioengineered BM confirms that the niche environment provides cytoprotection to cancer cells against chemotherapy^53^ (**Fig. 5 B&C**). Thirdly, our model (**Fig. 6 A&B**) confirms *in vivo* observations of elevated taurine metabolism in AML. We also show in human marrow samples that taurine metabolism increases further in AML patients who do not respond to AraC and decreases in responders (**Fig. 6 C&D**), and that GSE1159 data links elevated taurine metabolism to worse patient outcomes (**Fig. 6 E-G**). Finally, our model further validates *in vivo* findings^48^ that taurine transporter inhibition helps AraC kill diseased cells (**Fig. 7f**).

## Materials and Methods

See supplementary table I and II for reagent and antibody information.

### Preparation of materials and ECM interfaces

Plasma equipment was set up as per previous work ^28,29^. Samples (Glass coverslips, Nunc™ Thermanox™ coverslips or Commercial Corning® 96-well tissue culture plates (TCP)) were placed in the plasma chamber vertically to plasma flow. Then, samples were exposed to air plasma for 5 minutes at 50W of radio frequency (RF) incident power, followed by PEA polymerisation for 15 minutes at 50W of RF incident power. Samples were sterilised under ultraviolet (UV) light for 30 minutes before use.

For ECM protein coating, PBS solution containing human FN (20 μg/mL), murine LM (20 μg/mL), or the mixture of human FN (10 μg/mL) and murine LM (10 μg/mL), were coated onto pPEA modified samples for 1 hour, termed as FN, LM and FN+LM respectively. For GF adsorption, BMP2, NGF, and CXCL12 were added into PBS alone or in combination (final concentration 50 ng/mL) and coated onto ECM-coated pPEA substrates for 2 hours, followed by washing with PBS. TCP was used as a control.

### XPS analysis

Plasma polymerised samples were sent out to the National EPSRC Users’ Service (NEXUS) (found at: http://www.ncl.ac.uk/nexus/) for X-ray photoelectron spectra (XPS) analysis. A K-alpha XPS system (Thermo Fisher Scientific) equipped with a monochromatic Al K-alpha source was used to analyse each sample three times at a maximum beam size of 400 μm × 800 μm. Parameters were set up as follows: X-ray energy: 1486.68 eV; voltage: 12 kV; current: 3 mA; power: 36 W. Spectra analysis and curve fitting were performed using CasaXPS software.

### Atomic force microscopy analysis

After ECM coating onto TCP/pPEA surfaces, samples were washed with PBS three times, and finally with deionised water. Samples were then dried under a stream of nitrogen for AFM imaging. The JPK Nanowizard 4 was used for imaging and the JPK Data Processing software version 5 was used for image analysis. A pyramidal silicon tip (MPP-21 220, Bruker) was utilised for alternating contact mode imaging, with a resonance frequency of 75 kHz and stiffness of 3 N/m.

For the immunogold experiment, samples were fixed with fixation buffer (4% formaldehyde) for 10 minutes at room temperature, followed by incubation with primary antibody for 1 hour at room temperature. After being washed with 0.1% Tween 20/PBS, samples were incubated with gold-particle conjugated secondary antibody for 1 hour at room temperature. After that, the samples were fixed again with the fixation buffer for 20 minutes at room temperature and then washed with MilliQ water. Samples were dried gently with nitrogen gas before AFM imaging.

### Availability of functional domains in ECM

After ECM coating onto TCP/pPEA surfaces, samples were blocked with blocking buffer (1% BSA/PBS) for 30 minutes at room temperature, followed by primary antibody incubation for 1 hour at room temperature. After being washed with 0.1% Tween-20/PBS, samples were incubated with HRP-conjugated secondary antibodies for 1 hour. After that, samples were incubated with substrate solution for 20 minutes and the reaction stopped by adding stop solution. Absorbance was read at 450 nm using a plate reader.

### ELISA

For quantification of GF absorption onto ECM coated substrates, GF solutions from each substrate were aspirated and collected. GF original solution and GF aspirate were diluted 20X. For the quantification of CXCL12, the supernatant from different samples was collected, followed by concentration via a protein concentrator. The pooled supernatant was added to the sample chamber of the concentrator and centrifuged at 12000 g for 30 minutes at 4°C. The resulting solution was diluted to 100 µL. ELISA was then performed according to the manufacturer’s instructions. Briefly, standards and samples were added to the capture-antibody coated plates and incubated for 2 hours at room temperature. After incubation, the detection antibody was added and incubated for 2 hours at room temperature. Streptavidin-HRP was then added to plates for 20 minutes protected from light, followed by substrate solution for 20 minutes followed by stop solution. The plates were read using a plate reader with an absorptance wavelength at 450 nm and 570 nm. The standard curve was calculated using a four-parameter logistic curve fit. The GF concentration was calculated from the standard curve based on standards of known concentration.

### MSC/HSC isolation and culture

Human MSCs and CD34^+^ cells were isolated from the bone marrow aspirates of patients undergoing joint replacement surgery. This work is covered by the NHS GGC Biorepository ethics for the collection of surplus samples for research, rec 22/WS/020. For isolation of MSCs, the bone marrow samples were passed through the EASYstrainer (Greiner Bio-One #542070, core size 70 µm) and then gently added to 7.5 mL Ficoll-Paque Premium buffer (Sigma, #17-544-02). After centrifuging at 350 g for 45 minutes, the mononuclear cell interface layer was collected. Mononuclear cells were seeded into tissue culture flasks and incubated at 37°C, 5% CO_2,_ in D10; Dulbecco’s Modified Eagle Medium (DMEM, Sigma-Aldrich) supplemented with 10% Fetal Bovine Serum (FBS, Sigma-Aldrich), 1% sodium pyruvate (11 mg/mL, Sigma-Aldrich), 1% Gibco non-essential amino acids (NEAA, Thermo Fisher Scientific) and 0.1 mg/mL penicillin/streptomycin (Sigma-Aldrich). After 3-day culture, the non-adherent cells were removed. For experiments, MSCs were seeded at 3000/cm^2^ (passage 2-4) onto materials in D2 medium (DMEM with 2% FBS, 1% sodium pyruvate, 1% NEAA and 0.1 mg/mL penicillin/streptomycin). The medium was changed twice weekly. Experiments were repeated 3 times with MSCs from 3 patient donors.

For isolation of CD34^+^ cells, mononuclear cells were resuspended with CD34^+^ buffer (PBS with 0.5 mM EDTA and 5% human serum albumin (HSA)) and incubated with CD34^+^ microbeads (Miltenyi Biotec, #130-046-702) for 30 minutes at 4°C. Cell suspensions were transferred to LS Columns (Miltenyl Biotec, #130-042-401) and passed through the magnetic Quadromacs Separation Unit (Miltenyi Biotec, #130-090-976). Magnetically labelled CD34^+^ cells were then collected, counted and cryopreserved.

### THP1 culture

THP1 cell line (Sigma #88081201) was maintained and cultured in R10 media (RPMI 1640 supplied with 2 nM Glutamine, 10% FBS and 0.1 mg/mL penicillin/streptomycin) under a humidified atmosphere of 5% CO_2_ at 37°C. Cell concentration was maintained at 3×10^5^/mL, media changed twice weekly.

For the AML niche model, 5×10^3^ THP1 cells were seeded in 1:1 mix of D2 media and R2 media (RPMI 1640 supplied with 2 nM Glutamine, 2% FBS and 0.1 mg/mL penicillin/streptomycin). Cells were harvested after 3 days for further analysis.

For the chemotherapy treatment, THP1 only or within niches were treated with Cytarabine (AraC, #PHR1787) at the indicated concentration (1.13 μM) for 3 days, with the presence or absence of P4S inhibitor.

For CFSE staining, THP1 cells were cultured in 1:1 mix of D2 media and R2 media for 16hrs before they were stained with carboxyfluorescein succinimidyl ester (CFSE, #C34570). 15 million of THP1 cells were pelleted by centrifugation at 300g for 5mins and resuspended in PBS containing 1µM of CFSE. Cells were placed in a 37 °C incubator for 10mins, followed by deactivation of the free dye by adding 5X the volume of 1:1 mix of D2 media and R2 media that was originally used for the staining. After 5 minutes, the cells were pelleted at 300g for 5 minutes and then resuspended in the appropriate volumes of D2/R2 media with or without 100 µM P4S. 5×103 THP1 cells were seeded and their proliferation was assessed after 3 days.

### HSC culture

CD34^+^ cells were cultured in Iscove’s Modified Dulbecco Medium (IMDM) containing 20% Bovine serum albumin (BSA), Insulin and Transferrin (BIT) 9500 Serum Substitute (STEMCELL Technologies), 2 mM L-glutamine, 0.1 mg/mL penicillin/streptomycin supplemented with Flt3L (50 ng/mL), SCF (20 ng/mL) and TPO (25 ng/mL). CD34^+^ cells were thawed, counted and rested overnight in media at 1×10^6^/mL, then recounted and seeded into niche models at 5×10^4^/mL (0.1 mL/well of 96 well plates). For conditions containing gels, HSCs were seeded on top of the gel. Control wells were set up with media only (HSC only control). After 5 days cultured cells were harvested for further analysis. Experiments were repeated 3 times with HSCs from 3 patient donors.

### Immunofluorescence and imaging analysis

For cell morphology and focal adhesion analysis, MSCs were seeded onto FN/FN+LM/LM coated TCP/pPEA surfaces for 3 h, followed by actin (Alexa Fluor™ 488 phalloidin) and anti-vinculin staining. For quantification of the nuclear-to-cytoplasmic YAP ratio, MSCs were seeded onto FN/FN+LM/LM coated pPEA surface for 3 hours, followed by actin (Alexa Fluor™ 488 phalloidin) and anti-YAP staining. To block integrin function, cells were incubated with the inhibitory antibodies for 30 minutes before being seeded onto the ECM-coated substrates. A control rat IgG isotype antibody (#I-4000, 1:20, Vector Laboratories) was also used. To phenotype the MSCs in the niche models, MSCs were cultured within the niche models for 14 days, followed by nestin/VCAM/OPN staining (see supplementary table II). Immunofluorescence staining was carried out according to our previous paper^25,29^. The samples were fixed with fixation buffer for 15 minutes at 37°C, followed by permeabilisation buffer [10.3 g of sucrose,0.292 g of NaCl, 0.06 g of MgCl_2_, 0.476 g of HEPES buffer, 0.5 mL of Triton X, in 100 mL of PBS (pH 7.2)] for 4 minutes at 4°C. Then samples were blocked with 1% BSA/PBS for 30 minutes and incubated with primary antibodies overnight at 4C. On the second day, the samples were washed 3x with 0.1% Tween-20/PBS and subsequently incubated with fluorophore-conjugated secondary antibody for 2 hours at room temperature. Before image acquisition, samples were incubated with DAPI/PBS buffer for 15 minutes. For quantification of cell number, cells were stained with Hoescht and nuclei counted. The images were acquired by EVOS M700 microscope (Thermo Fisher Scientific).

Vinculin images were used for FA analysis ^54^. Images were analysed with a threshold area of 0.5 µm^2^ and 0–0.99 circularity. Binarized images of FAs were measured, and several parameters were used for FA analysis, including FA number per cell, the FA length and FA area.

The mean integrated fluorescent intensity (MFI) was determined by ImageJ. For YAP translocation analysis, the total intensity of YAP was divided by the total intensity of YAP in the cytoplasm to obtain the nuclear-to-cytoplasmic ratio. At least 10 cells were analysed per donor per group.

### Western blotting

MSCs were seeded on FN/FN+LM/LM coated pPEA surfaces for 1 hour, protein was extracted using lysis buffer containing protease and phosphatase inhibitors. Protein concentrations were measured using the BCA Protein Assay Kit. Heat-denatured proteins were electrophoresed by 4-12% Bis-Tris Gel and then transferred onto PVDF membranes. The membranes were blocked in 5% milk/PBS for 30 minutes at room temperature and incubated with primary antibodies (pFAK and MLC) overnight at 4°C. Samples were then incubated with the appropriate HRP-conjugated secondary antibody for 2 hours at room temperature. Membranes were read using enhanced chemiluminescence reagents (Thermo Fisher Scientific, #32132) and visualised using a detection system (Thermo Fisher Scientific MYECL imager). Band intensities were analysed using Image J software and the intensity fold change over the FN group was calculated for statistical analysis.

### In-cell Western

To quantify ECM absorption onto pPEA surface, ECM-coated TCP/pPEA surfaces were washed 3x PBS and stained using FN or LM antibodies (see supplementary table II). For quantification of expression levels of nestin, VCAM, and OPN in MSCs on ECM+GFs surfaces, MSCs were cultured on the indicated surfaces for 14 days,. In-Cell Western was carried out according to previous work^25,29^. Samples were fixed with fixation buffer for 15 minutes at 37°C, followed by permeabilisation buffer for 4 minutes at 4°C. Then samples were blocked with 1% milk/PBS for 1.5 hours and incubated with primary antibody overnight. Then, samples were washed with the 0.1% Tween-20/PBS buffer x5 and incubated with infrared-labelled secondary antibodies for 2 hours, and washed x5. Samples were dried in the fridge overnight, and scanned using a LiCor Odyssey infrared imaging system. Samples were stained with CellTag 700 to detect cell number and used to normalise against the protein of interest.

### Cytokine array

After 14 days of culture, supernatant from MSCs seeded on FN/FN+LM/LM coated pPEA surfaces was collected for the cytokine array. Membranes were blocked for 1 hour and incubated with samples of desired dilution overnight at 4°C. membranes were then washed and incubated with detection antibody for 1 hour. Membranes were then again washed, and incubated with streptavidin-HRP. Chemi Reagent Mix was added and spread evenly across the membranes for imaging. Odyssey infrared imaging system was used to quantify the signal intensity. Intensity fold changes over the FN group were calculated for analysis.

### AlamarBlue assay

5×10^3^ THP1 cells were seeded in 96 well plates and treated with AraC at serial concentrations (from 0 to 1000 uM). After 72-hour treatment, 10 µl of AlamarBlue resazurin was added to each well, followed by further incubation for 6 hours at 37°C, 5% CO_2_. A microplate reader (Clariostar, Germany) was used to detect light adsorbance at wavelengths of 570 nm and 600 nm. Non-treated THP1 was used as the control. The percentage of AlamarBlue reduction to control was calculated and the IC_50_ determined as the fifty-percent point intercepting the Dose-Response Curve to the concentration along the x-axis.

### FN PEGylation

FN was covalently connected with 4-arm-poly (ethylene glycol) (PEG)-maleimide (PEG-MAL) molecules on its thiol groups. This process of PEGylation has been demonstrated in our previous work ^39^ and others’ work^55^. FN was incubated within denaturing buffer with 8 M urea, 5 mM TCEP (Tris (2-carboxyethyl) phosphine) in PBS, followed by the incubation with 4-arm-PEG-maleimide solution at a molar ratio of 4:1 (PEG-MAL: FN) for 30 minutes. The reaction was stopped by adding 0.5 µl NaOH (1 M stock). Subsequently, the mixture was incubated with 14 mM of iodoacetamide for 2 hours at room temperature, followed by dialysis against PBS for 1 hour. Then, the protein solution was incubated with nine volumes of absolute ethanol at -20°C overnight. Then, the solution was centrifuged at 15000 g for 15 minutes at 4°C and washed with absolute ethanol twice. After the final wash, the protein precipitation was air dried for 30 minutes at room temperature and then dissolved in 8 M urea producing a protein concentration of 2.5 mg/mL, followed by dialysis again against PBS for 1 hour at room temperature. The molecular weight cut-off of the dialysis tube was 10 kDa. The final dialysed protein was then stored at -20°C until use.

### Hydrogel preparation

PEG-MAL was diluted by PBS to 200 mg/ml. The crosslinkers, PEG-Dithiol (PEG-SH) or protease-degradable VPM peptide (GCRD**VPM**SMRGGDRCG, Genscript), were diluted in PBS to 50 mg/ml. Then, PEG-4MAL and crosslinkers were mixed at a molar ratio of 1:1 (maleimide: thiol) to crosslink. Crosslinkers used were mixtures of PEG-SH and VPM at a molar ratio of 1:3. The final concentration of PEG-4MAL is 3% w/v. Following addition of crosslinkers, samples were incubated for 30 minutes at 37°C to for gelation. For FN-PEG hydrogel, PEGylated FN was added at a final concentration of 0.5 mg/ml. For hydrogels with labelled GFs, the labelled GFs were mixed into the hydrogel solution with a final concentration of 5 μg/mL prior to gelation.

### Nanoindentation measurement and data analysis

Nanoindentation measurements were performed using a nanoindentation device (Chiaro, Optics11) mounted on top of an inverted phase-contrast microscope (Evos XL Core, Thermo Fisher Scientific) adapting a previously reported approach ^56,57^. Measurements were performed at room temperature in culture media unless stated otherwise. For each hydrogel sample, single indentation curves (n > 22) were acquired at a speed of 2 µm/s over a vertical range of 10 µm, changing the (x, y) point at every indentation. The selected cantilever had a stiffness of 0.032 N/m and held a spherical tip of 8.0 µm radius. For cell stiffness measurements, cells were cultured on FN or LM pPEA substrates for 3 hours to allow for cell adhesion to the surfaces. For each cell, indentation maps (400 µm^2^, n > 25) were also acquired at a speed of 2 µm/s over a vertical range of 10 µm, changing the (x, y) point at every indentation. A minimum of three maps per replicate were measured. The selected cantilever had a stiffness of 0.36 N/m and held a spherical tip of 22 µm radius. All collected curves were pre-processed and analysed with a custom-made graphical user interface, the analysis and all software used are described in detail in ^57^.

### GF labelling

GFs were conjugated with NHS-AlexaFluor-488 dye according to the manufacturer’s instructions. In brief, recombinant GF was dissolved with 0.05 M sodium borate buffer at pH 8.5 to 100 ug/mL. NHS-AlexaFluor-488 dye was mixed with the GF solution and incubated for 1 hour at room temperature, followed by dialysing against PBS for 4 hours. The molecular weight cut-off of the dialysis tube was 3.5 kDa. The final dialysed labelled GF was stored at - 20°C until use.

### GF absorption experiment

Prepared hydrogels were immersed in 10 mM L-cysteine solution for 2 hours to ensure no free maleimide groups. Then, samples were incubated within PBS overnight to swell. Then, hydrogels were immersed in solutions containing different labelled GFs at 5 μg/mL. Once immersed, samples were incubated overnight at 37°C protected from light. The supernatant was taken and read using a plate reader (Ex/Em 493/518 nm). All conditions were prepared in triplicate. The initial solutions were also measured and used as a standard curve to be able to correlate fluorescence intensity with the concentration of specific GFs. The percentage of GF adsorbed by each hydrogel was calculated by the following equation:

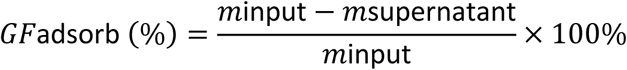

### GF retention experiment

Hydrogels with labelled GFs were immersed in PBS and incubated at 37°C protected from light. At each time point, all the supernatant was collected and refreshed with PBS. The GF concentration in the supernatant was determined by using a plate reader (Ex/Em 493/518 nm). A standard curve using the specific GF was prepared and empty (not loaded with GF) hydrogels were used as controls. All conditions were prepared in triplicate. The percentage of GF retained from each hydrogel was calculated by the following equation:

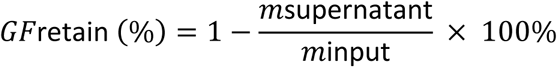

### Flow cytometry

Cells were harvested from niche systems using collagenase D (Sigma, 2.5 mg/mL in PBS) and TripLE™ (Thermo Fisher Scientific). Cells were passed through a 70 µm filter and stained with the antibodies detailed in **Table S2** for 30-45 minutes. Cells were washed twice in flow buffer (PBS with 2 mM EDTA and 0.5% BSA) and analysed using an Attune flow cytometer (Thermo Fisher Scientific). Unstained cells or isotype controls were used as a negative control. To phenotype HSCs, the gating strategy is shown in **Fig. S9A**. For LTC-IC assays, samples were sorted using BD FACSAria Cell sorter (BD Biosciences). The gating strategy is shown in **Fig. S10**. For mtROS staining, cells were stained with MitoSOX probe for 10 minutes at 37°C, and then washed with HBSS buffer 3 times followed by flow cytometry analysis. The gating strategy is shown in **Fig. S13A**. For PI staining, cells were stained with PI dye for 15 minutes at room temperature, and then samples were added with 400 μl flow buffer. Fixed THP1 were used as a positive control. For Zombie violet staining, cells were stained with Zombie Violet stain at 1:500 dilution on ice for 15 minutes (Biolgened, #423113). 1ml of flow buffer was added to deactivate the leftover dye and the sample viability was assessed by flow cytometry. Flow cytometry files were analysed using FlowJo software (version 10.5.3, FlowJo LLC, USA) or FCS Express v7.

### LTC-IC assay

LTC-IC assays were carried out on sorted Lineage^-^ CD34^+^ cells harvested from niche systems after 5 days of culture. Engineered stromal fibroblast feeder layers were first established. M2-10B4 (overexpressing IL-3 and G-CSF) and Sl/Sl (overexpressing IL-3 and SCF) (both gifted by STEMCELL Technologies) were passaged at ∼90% confluence. Cells were grown for 2 weeks prior to use in selection agents to select stromal cells expressing long-term cell maintenance factors (M2-10B4, 0.4mg/mL G418 and 0.06mg/mL Hygromycin B; Sl/Sl, 0.8 mg/mL G418 and 0.15 mg/mL Hygromycin B). Cells at ∼80% confluence were irradiated with 8000 cGy, and trypsinised. M2-10B4 and Sl/Sl were mixed at 1:1 ratio at a final concentration of 1.5×10^6^/mL. Cells were then seeded into collagen-coated 24 well plates (Thermo Fischer Scientific) for 24 hours before adding sorted cells. The culture was then maintained for 6 weeks in MyeloCult™ supplemented with 1×10^-6^ M hydrocortisone (StemCell Technologies), with half media exchanges twice weekly.

### CFU Assay

Cells were harvested from LTC-IC assays after 6 weeks and resuspended in MethoCult™. 3×10^5^ cells for each condition were seeded in duplicate in 35 mm dishes and incubated at 37°C in 5% CO_2_. After 14 days, total colonies were counted and phenotyped using a light microscope.

### RNA seq and data analysis

For RNA seq analysis of MSCs, MSCs were harvested from 3D niche model, healthy niche model and AML niche model after 3 days of co-culture (**Fig. 5A**). Sequencing libraries were then prepared from total RNA using the Illumina TruSeq Stranded mRNA Sample Preparation Kit. Libraries were sequenced in 75 base, paired-end mode on the Illumina NextSeq 500 platform. Raw sequence reads were trimmed for contaminating sequence adapters and poor-quality bases, followed by “pseudo aligned” to the transcriptome using the program Kallisto^58^. Human ENSEMBL gene ID to gene symbol conversion was performed in BioTools (https://www.biotools.fr). The DEseq2 BioConductor package was used for outlier detection, normalisation and differential gene expression analyses ^59^. Genes passing a threshold of adjusted p-value <0.05 and |log2Foldchange| > 1 were considered as differentially expressed genes (DEGs). Heatmaps of DEGs, and principal component analysis (PCA) plots were generated in R. The differentially expressed genes along with their respective log2fold change were inputted into the PathfindR package, followed by Kyoto Encyclopedia of Genes and Genomes (KEGG) and Gene Ontology (GO) enrichment analysis^60^. Only pathways with FDR < = 0.05 were considered as differentially enriched. The differentially expressed genes involved in every pathway were extracted and their z-scores were presented as heatmaps.

### Metabolomics and data analysis

MSCs were harvested from niche models after 3 days of co-culture (**Fig. 5A**). Whole-cell metabolomic analysis was performed on cell lysates isolated from MSCs cultured in niche systems. Substrates were washed with ice-cold PBS, and cells lysed in extraction buffer (PBS/methanol/chloroform at 1:3:1 ratio) for 60 minutes at 4°C with constant agitation. Lysates were then transferred to cold Eppendorf tubes and centrifuged at 13000 g at 4°C for 5 minutes to remove debris and stored at -80°C. Cleared extracts were used for hydrophilic interaction LC/MS analysis (UltiMate 3000 RSLC, Thermo Fisher Scientific), with a 159 x 4.6 mm ZIC-pHILIC column running at 300 µl/min and Orbitrap Exactive (Thermo Fisher Scientific). A standard pipeline, consisting of XCMS (peak picking), MzMatch (filtering and grouping) and IDEOM (further filtering, post-processing and identification) was used to process the raw mass spectrometry data. Identified core metabolites were validated against a panel of unambiguous standards by mass and predicted retention time. Further, putative identifications were generated by mass and predicted retention times. Heatmaps of selected metabolites, PCA plots, and enrichment analysis were generated using MetaboAnalyst software (version 4.0).

### Immunohistochemistry

A total of 24 paraffin-embedded bone marrow samples at Peking Union Medical College Hospital, Beijing, China, were included in our current study. Among them, 12 patients were diagnosed with AML by experienced pathologists, termed AML donors. The other 12 patients’ bone marrow samples were collected because of accident fractures, termed healthy donors. The paraffin-embedded samples were cut into 4-μm-thick sections, followed by staining using Nestin antibody (ab22035; 1:2000; Abcam). Afterwards, the slides were incubated with secondary antibody at room temperature for one hour and stained with DAB substrate. All specimens were stained using an automatic IHC instrument (BOND-III; Leica Biosystems), according to the manufacturer’s instructions.

For immunohistochemistry evaluation, all slides were scanned using a Hamamatsu S60 whole slide scanner (Hamamatsu Photonics), and the digitally scanned images were viewed and evaluated using NanoZoomer Digital Pathology view2 software version 2.7.25 (Hamamatsu Photonics). The nestin-postive area was quantified by IHC Profiler plug-in in ImageJ software^61^.

### Statistical analysis and reproducibility

Data were presented as means ± standard deviation (s.d.). The unpaired Student’s t-test was applied for two-group comparisons except for fig. 4b, fig. 5b-c, fig 6d, fig. 7d-f, supplementary fig. 8a and 8b (paired Student’s t-test). The ordinary two-way ANOVA with Tukey’s multiple comparisons test was applied for multi-group comparisons, except for fig. 3E and 3F (Ordinary two-way ANOVA with Dunnett’s multiple comparisons test). GraphPad Prism 8TM software was used for statistical analysis. All the experiments were repeated with 3-4 material replicates for 3 independent experiments with different donor cells, except for cytokine array (1 independent donor with technical duplicates) and LTC-IC assay (2 independent donors with >3 technical repeats).

## Acknowledgments

We thank Carol-Anne Smith and Dr Alasdair McDonald for technical assistance. We thank Glasgow Polyomics Facility for support with RNASeq and metabolomics and we thank Glasgow Imaging Facility, particularly Dr Leandro Lemgruber. This work was supported by a CSC scholarship [201908060063], Wellcome ISSF [204820/Z/16/Z] and National Natural Science Foundation of China [82300266] to X.Y, BBSRC and EPSRC funded grants BB/N018419/1, EP/P001114/1 and EP/X036049/1 to M.D and M.S-S. This study was approved by the Ethics Review Committee of Peking Union Medical College Hospital (Approval Number: K-6198).

## Author contributions

YX, SI, HD, ER, MS-S and MJD conceived and designed the analysis. XY, SI, HD, XL, MAGO and MPT performed the experimental work. OD, ST, MS, VJ and MV helped produce and characterise the surfaces and hydrogels. RMDM and PSY provided bone marrow for cell isolation. MC provided advice and guidance on LT-HSCs and chemotherapeutics. XY and MJD wrote the manuscript. YX, SI, HD, XL, OD, ST, MS, MAGO, PSY, MPT, VJ, MV, ER, MC, RMDM and MS-S revised the manuscript and were involved in the discussion of the work. XY, MS-S and MJD secured funding.

## Data and materials availability

Source data are provided with this paper. All data supporting the findings in this study are available within the article and its Supplementary Information files, can be obtained from the corresponding author or can be accessed at: https://doi.org/10.5525/gla.researchdata.1676

## Supplementary data

**Fig. S1.**
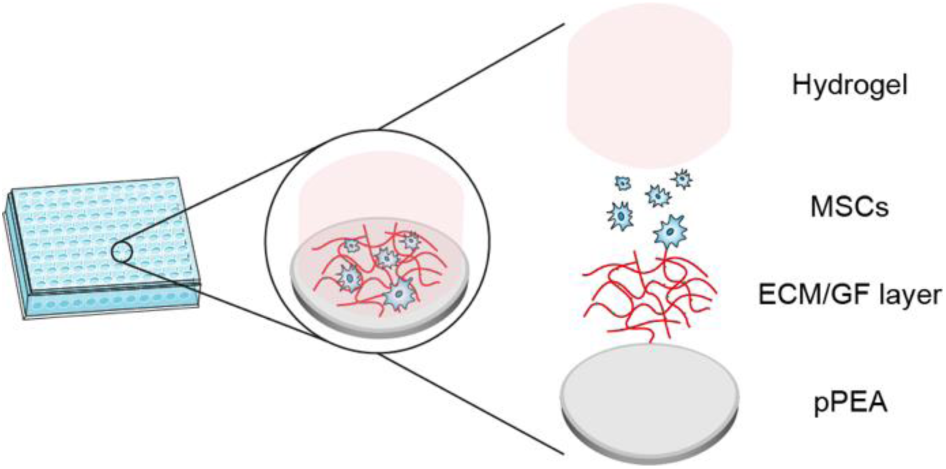
Schematic of the bone marrow niche model. PEA was plasma polymerized onto 96-well plates. Then, the blend of ECM proteins was coated on the pPEA surface, which can be followed by growth factor (GF) tethering, to mimic the in vivo BM ECM and GF microenvironment. As the key niche cellular population, human MSCs were then cultured within this system and directed into the HSC support phenotype. To mimic the stiffness and mechanics of the BM, PEG-FN hydrogels were next added on the top. Such a model provides a high-throughput platform to investigate the role of niche aspects in MSC behaviours, such as supportive capacity to HSC self-renewal and to leukemic progression during leukemic remodelling.

**Fig. S2.**
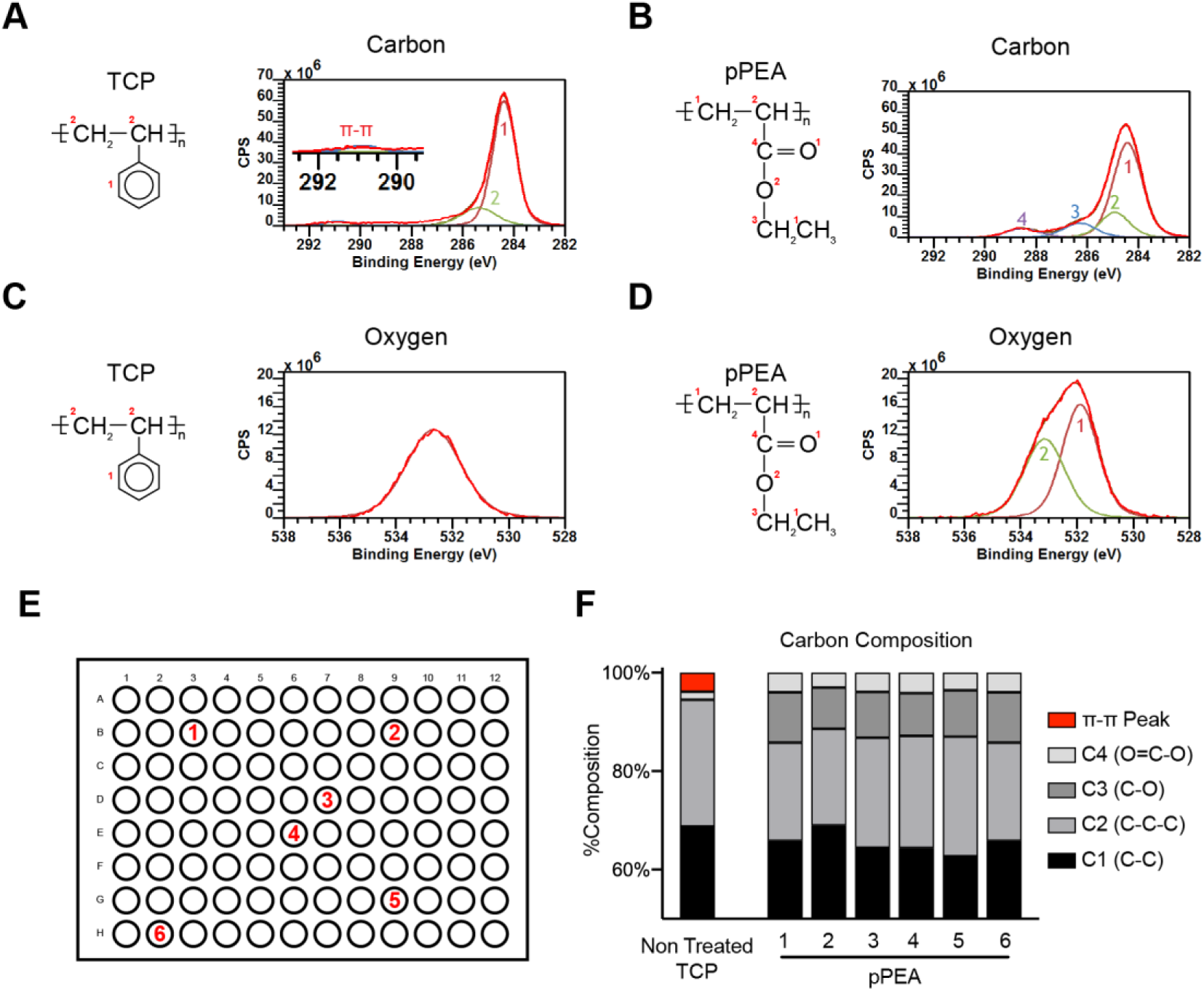
PS analysis of the surfaces. Carbon analysis shows that TCP (polystyrene) contains an aromatic group, and so a π− π bonding peak is observed on the carbon spectra at ∼291 eV (**A**). This feature is lost with the PEA coating. We instead observed peaks corresponding to features of the PEA polymer, particularly the ester group, indicative of successful coating (**B**). Oxygen spectrum for XPS analysis of TCP **(C)** and pPEA **(D)** surfaces. Chemical structures of polystyrene (TCP) and PEA are shown. Peaks were numerically labelled, corresponding to the representative chemical structure insets. PEA was polymerised evenly across 96-well plates. **E,** Schematic of randomly selected wells in the 96-well plate. **F,** The bar plot representing the calculated percentage of carbon composition in each well, demonstrating pPEA coating.

**Fig. S3.**
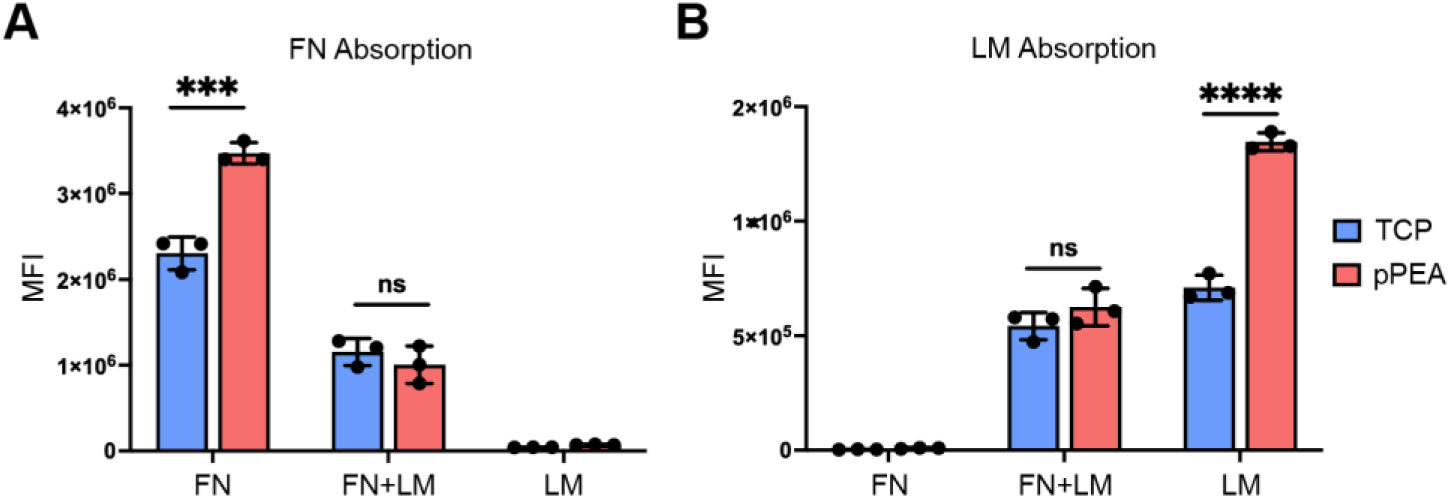
FN and LM absorption on TCP and pPEA surfaces. **A & B,** FN **(A)** and LM **(B)** absorption on TCP and pPEA substrates were quantified by in-cell western analysis. Mean ± s.d. based on n= 3 material replicates. ***, p<0.001, ****, p<0.0001, ns, non-significant. P values were determined by unpaired Student’s t-test.

**Fig. S4.**
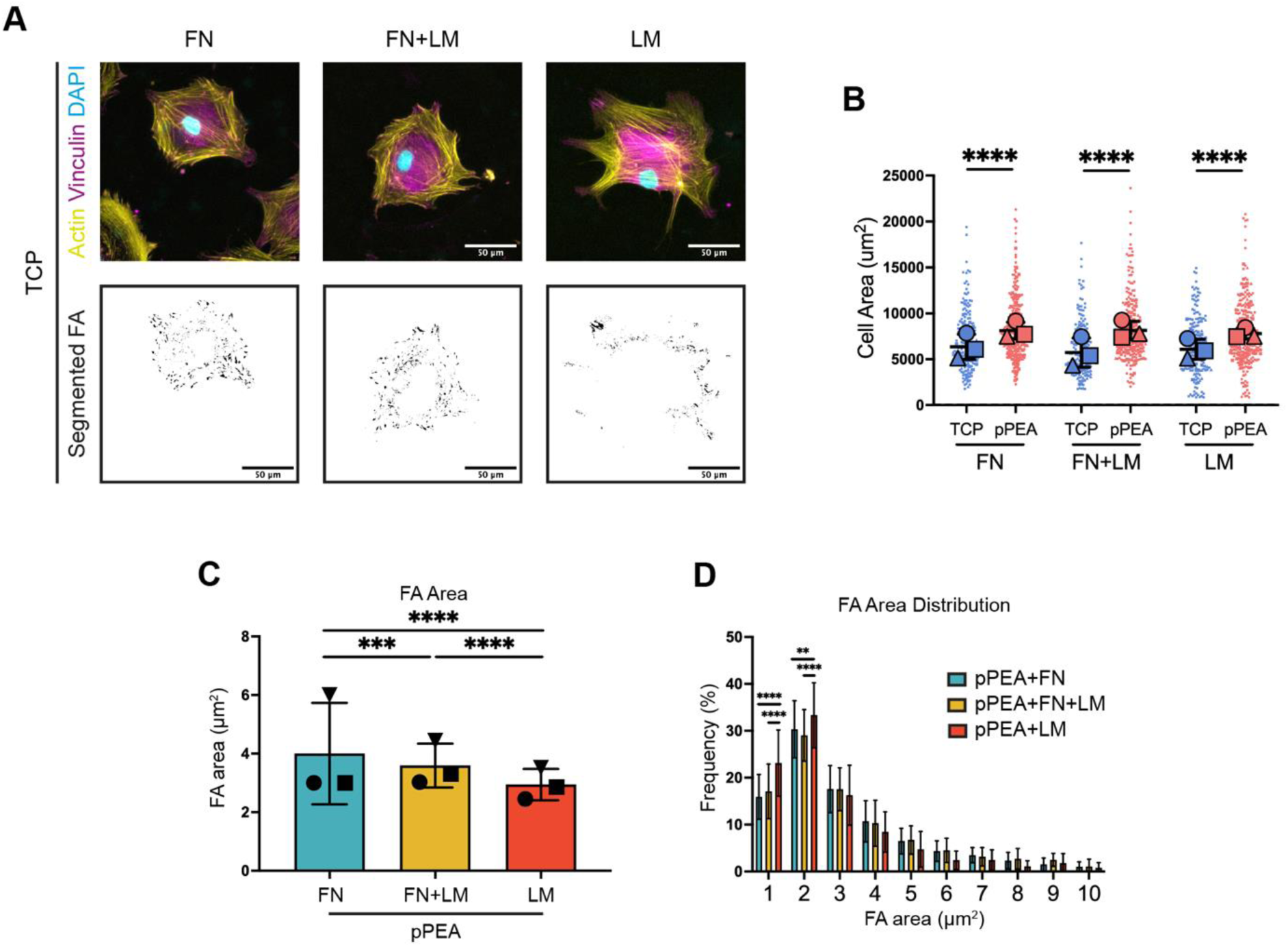
Cell morphology and adhesion area analysis of MSCs on FN, FN+LM, and LM coated surfaces. **A,** Representative immunofluorescence images of vinculin (magenta)/ actin (yellow)/ DAPI (blue) staining and thresholded binary images for MSC focal adhesion (FA) analysis. Scale bar: 50 μm. **B,** Cell area analysis of MSCs cultured on FN, FN+LM, and LM-coated TCP/pPEA surfaces. Each shape represents each donor, each small shape represents each cell, and each large shape represented the mean from each donor. Mean ± s.d. based on n=3 independent experiments with different donor cells. ****, p<0.0001. P values were determined by unpaired Student’s t-test. Analysis of FA area **(C)**, and FA area distribution **(D)** based on the vinculin/actin/DAPI staining. Each shape represents each donor. Mean ± s.d. based on n= 3 independent experiments with different donor cells. **, p<0.01, ***, p<0.001, ****, p<0.0001, ns, non-significant. P values were determined by ordinary two-way ANOVA with Tukey’s multiple comparisons test.

**Fig. S5.**
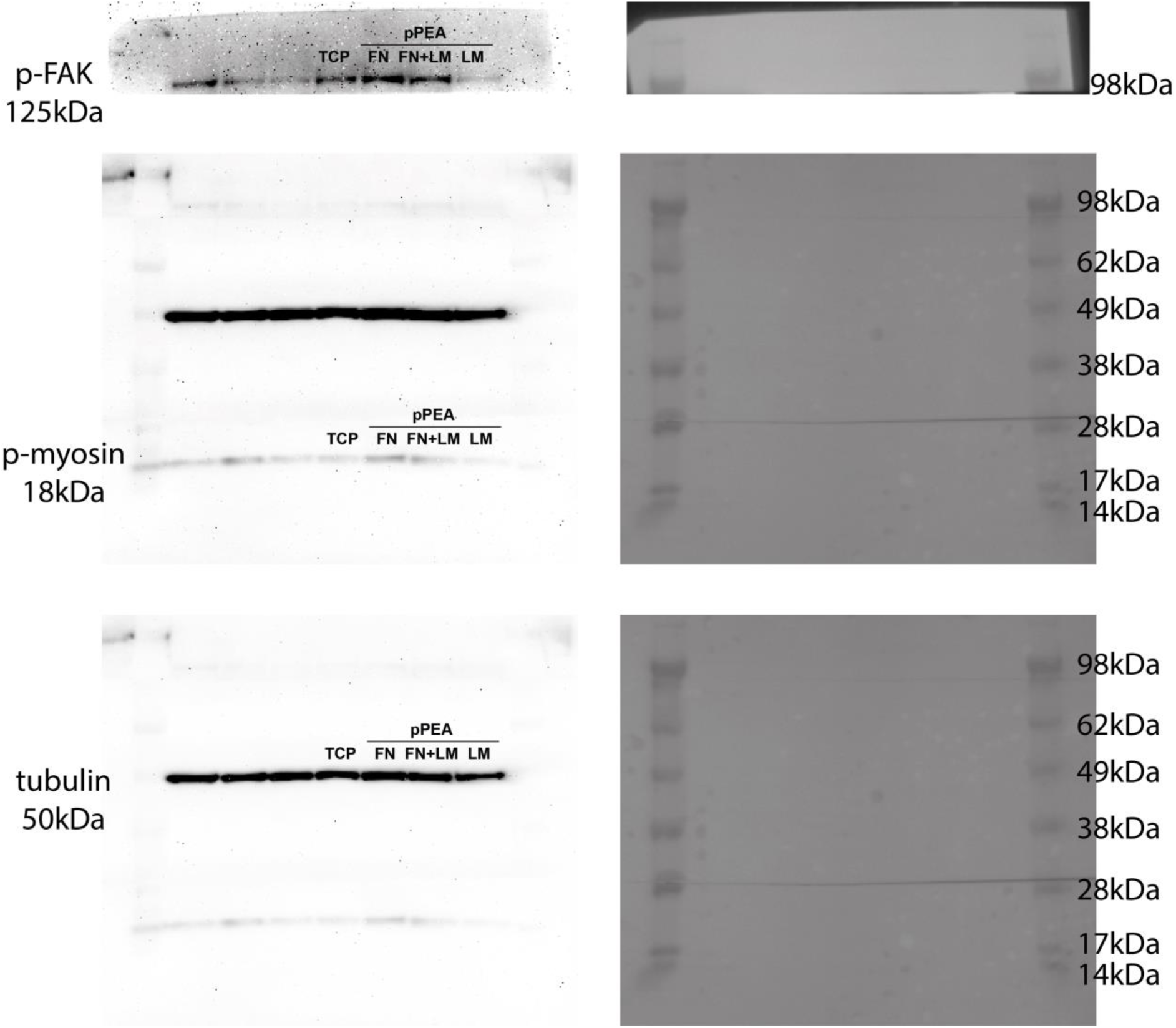
The full western blot as shown in Fig. 3F.

**Fig. S6.**
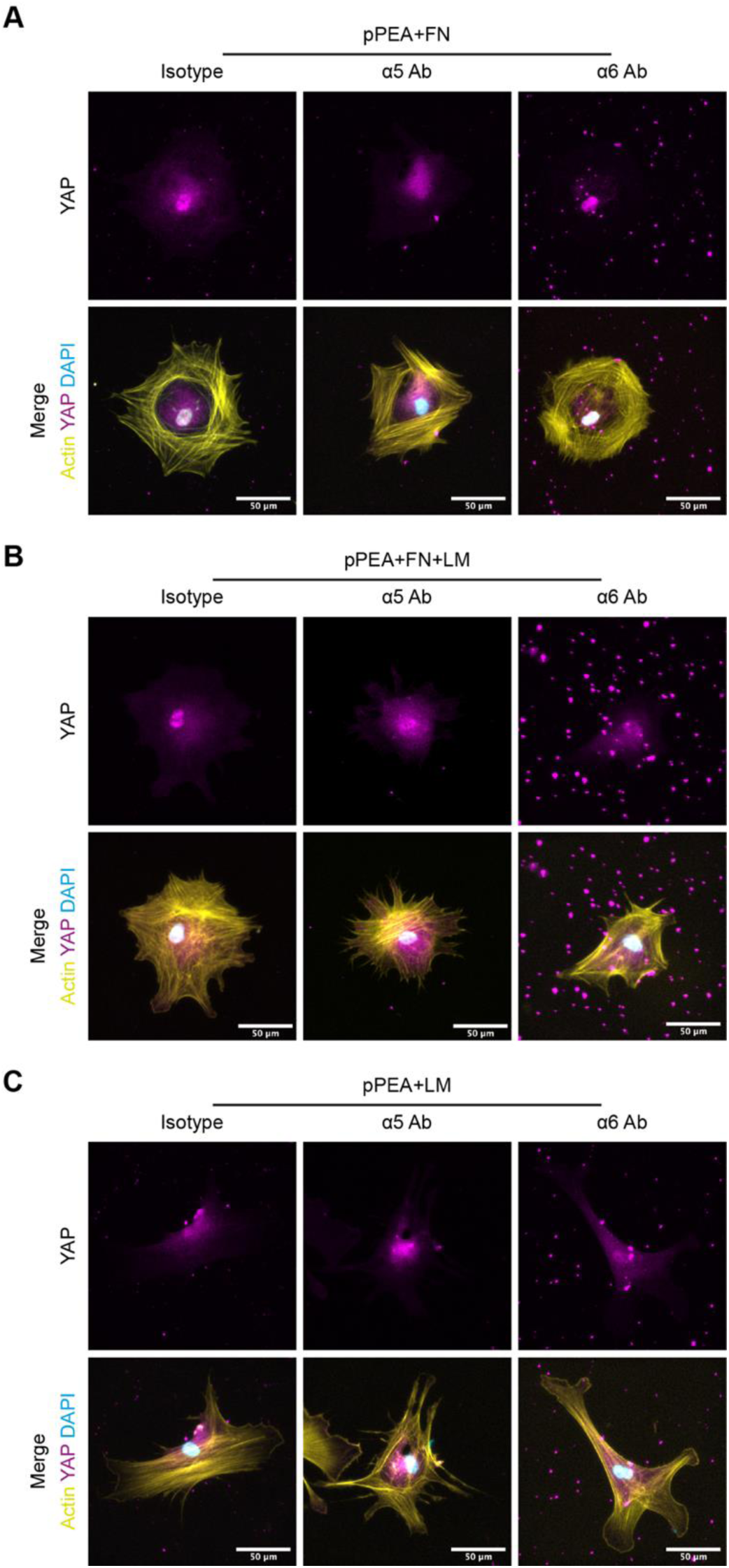
Representative images of YAP staining. **A,** Representative images for fig 3J-L of YAP (magenta)/ actin (yellow)/ DAPI (blue nuclear stain) staining in MSCs cultured on the pPEA+FN surface after α5/α6 integrin blocking. **B,** Representative images of YAP (magenta)/ actin (yellow)/ DAPI (blue) staining in MSCs cultured on the pPEA+FN/LM surface after α5/α6 integrin blocking. **C,** Representative images of YAP (magenta)/ actin (yellow)/ DAPI (blue) staining in MSCs cultured on the pPEA+LM surface after α5/α6 integrin blocking. Scale bar: 50 μm.

**Fig. S7.**
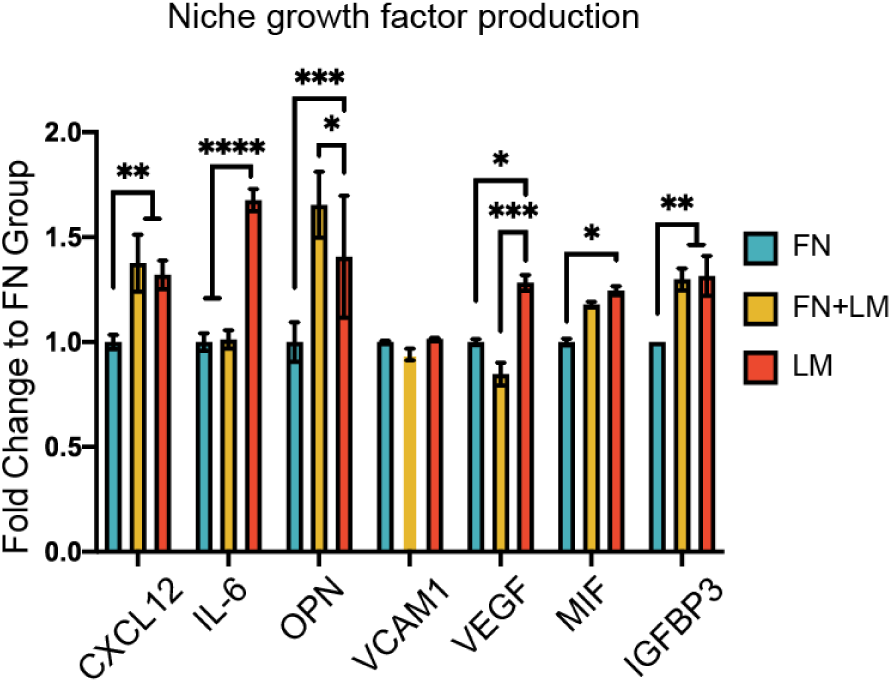
LM induces MSC to secrete niche-related GFs. Niche-related GF production for MSCs on FN, FN/LM, LM pPEA surfaces was quantified by cytokine array. Mean ± s.d. based on 1 independent donor with technical duplicates. *, p< 0.05, **, p< 0.01. ***, p<0.001, ****, p<0.0001, ns, non-significant. P values were determined by ordinary two-way ANOVA with Tukey’s multiple comparisons test.

**Fig. S8.**
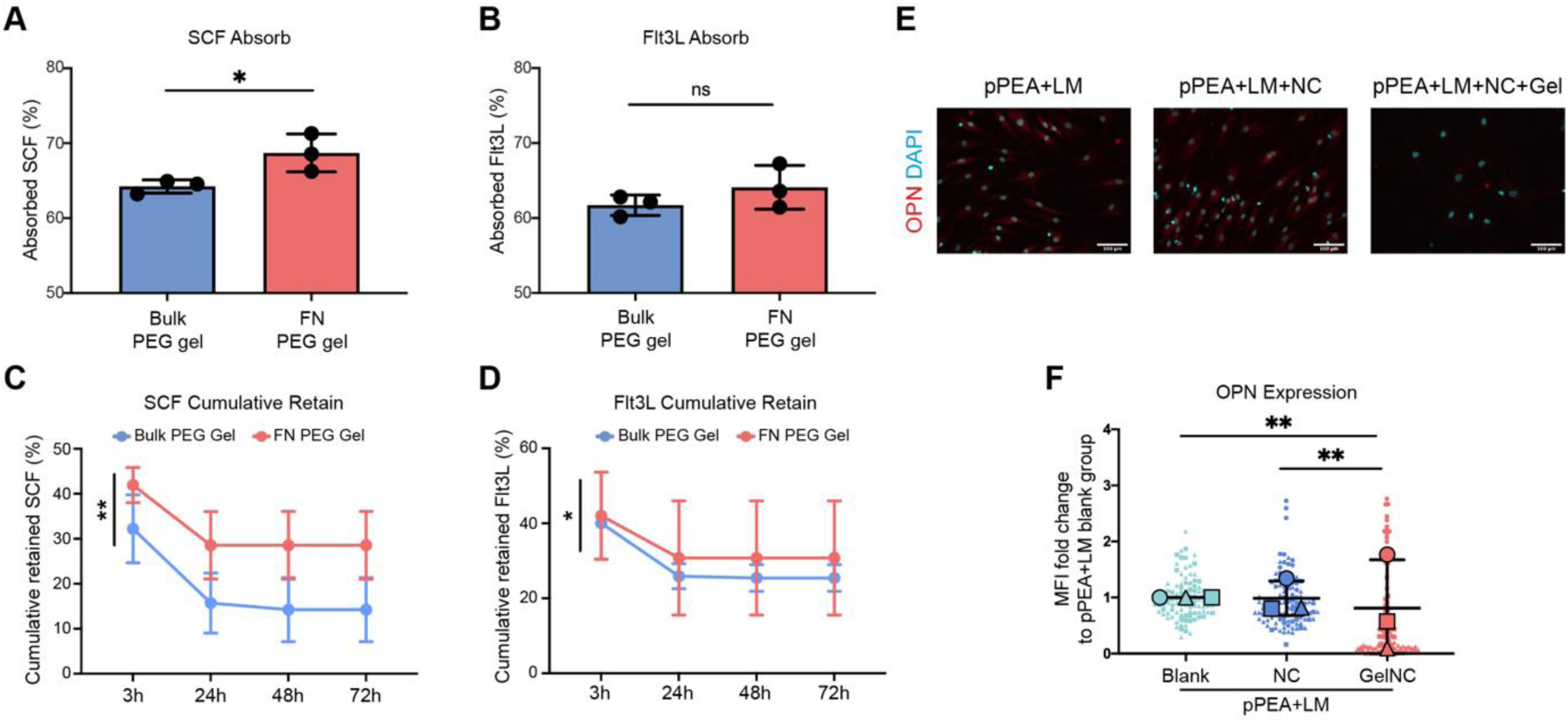
Absorption and retention of GFs in the PEG hydrogels and OPN expression of MSCs in the gels. **A & B**, The percentage of adsorbed SCF and Flt3L in bulk PEG gel and FN PEG gels. **C & D**, The percentage of cumulatively retained SCF and Flt3L in bulk PEG gel and FN PEG gels. Mean ± s.d. based on n= 3 material replicates. *, p<0.05, **, p<0.01, ns, non-significant. P values were determined by paired Student’s t-test (**A, B**) and unpaired Student’s t-test (**C, D**). **E,** Representative images of OPN staining in MSCs within the niche. Scale bar: 100μm. **F,** Immunofluorescence of OPN expression of MSCs cultured in the NC-containing models with (GelNC) /without (NC) FN-PEG gels for 14 days. Quantification analysis of MFI fold change values over the pPEA/LM (Blank) group is presented. Each shape represents each donor, each small shape represents each cell, and each large shape represents the mean from each donor. Mean ± s.d. based on n= 3 independent experiments with different donor cells. *, p<0.05, **, p<0.01, ***, p<0.001, ****, p<0.0001. P values were determined by ordinary two-way ANOVA with Tukey’s multiple comparisons test. (NC: NGF/CXCL12).

**Fig. S9.**
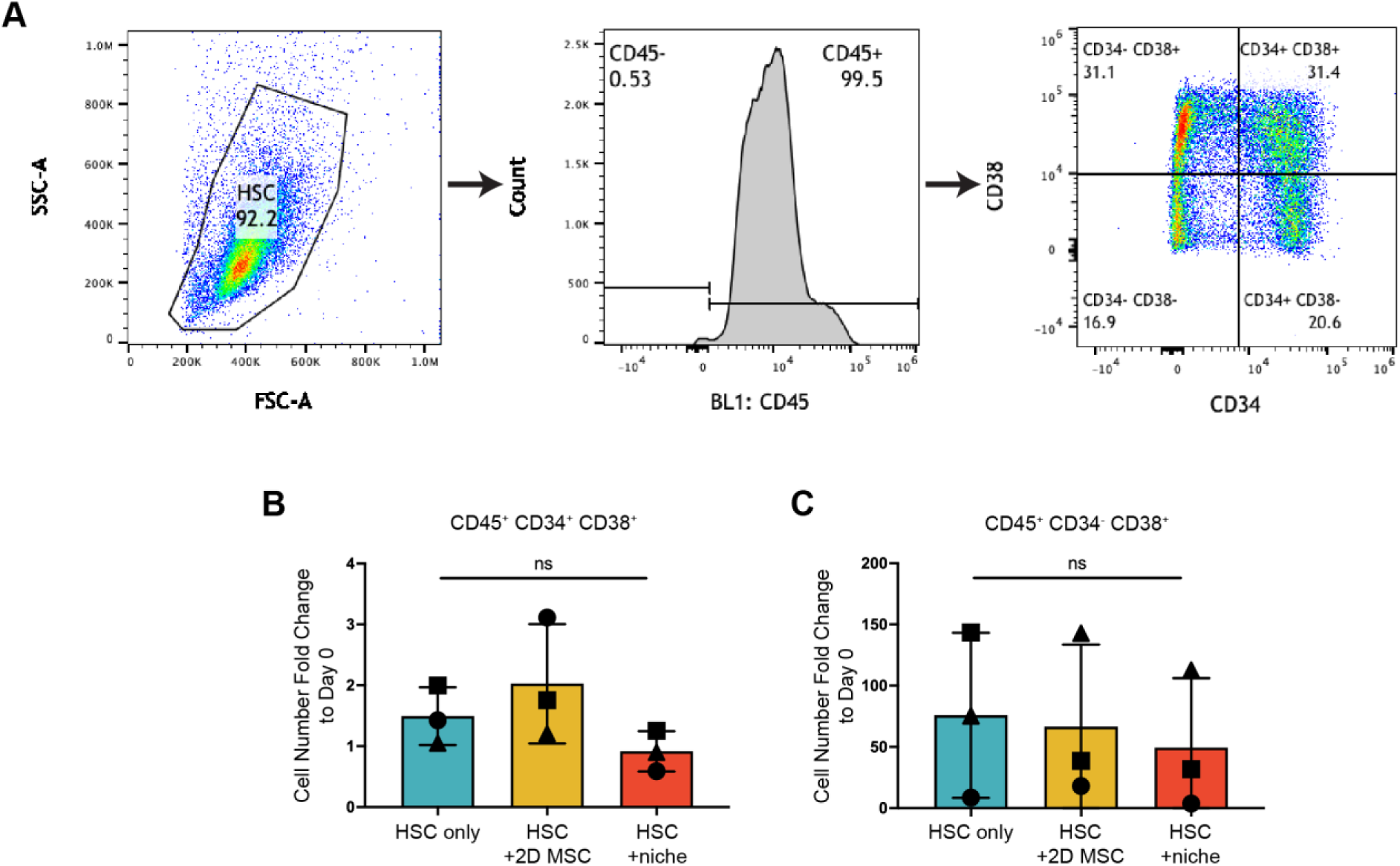
HSC phenotype in the niche model. **A,** FACS gating strategy for HSC and progenitor identification. 3 gates were used to identify CD34^+^CD38^-^ (LT- and ST- HSCs), CD34^+^CD38^-^ (HSPCs) and CD34^-^CD38^+^ (committed progenitors). **B & C,** Number of HSPCs (CD45^+^CD34^+^CD38^+^) and committed progenitors (CD45^+^CD34^-^CD38^+^) cultured in different niche models for 5 days. Each shape represents each donor. Mean ± s.d. based on n= 3 independent experiments with different donor cells. ns, non-significant. P values were determined by ordinary two-way ANOVA with Tukey’s multiple comparisons test.

**Fig. S10.**
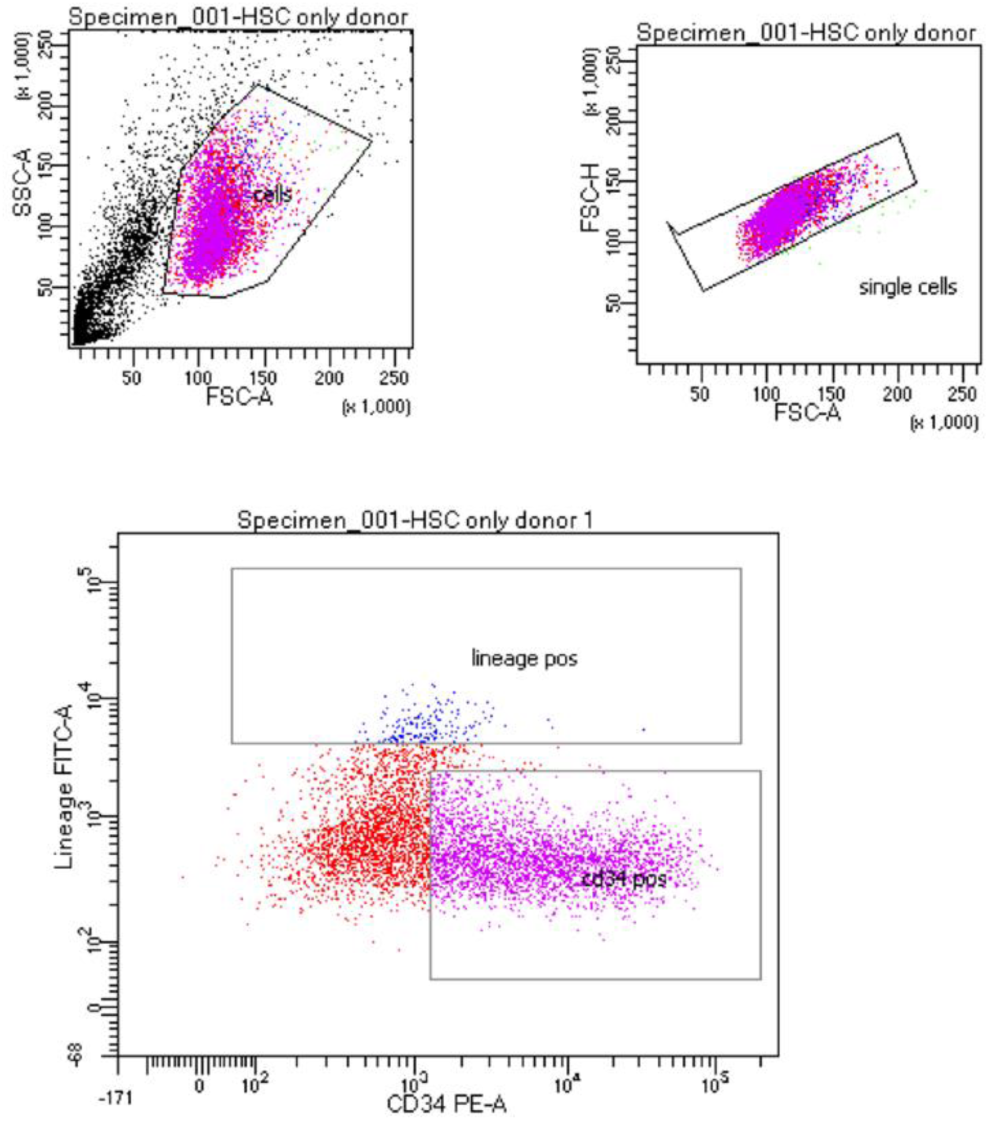
FACS gating strategy for cell sorting for LTC-IC. Lin^-^CD34^+^ cells were gated and sorted for LTC-IC analysis.

**Fig. S11.**
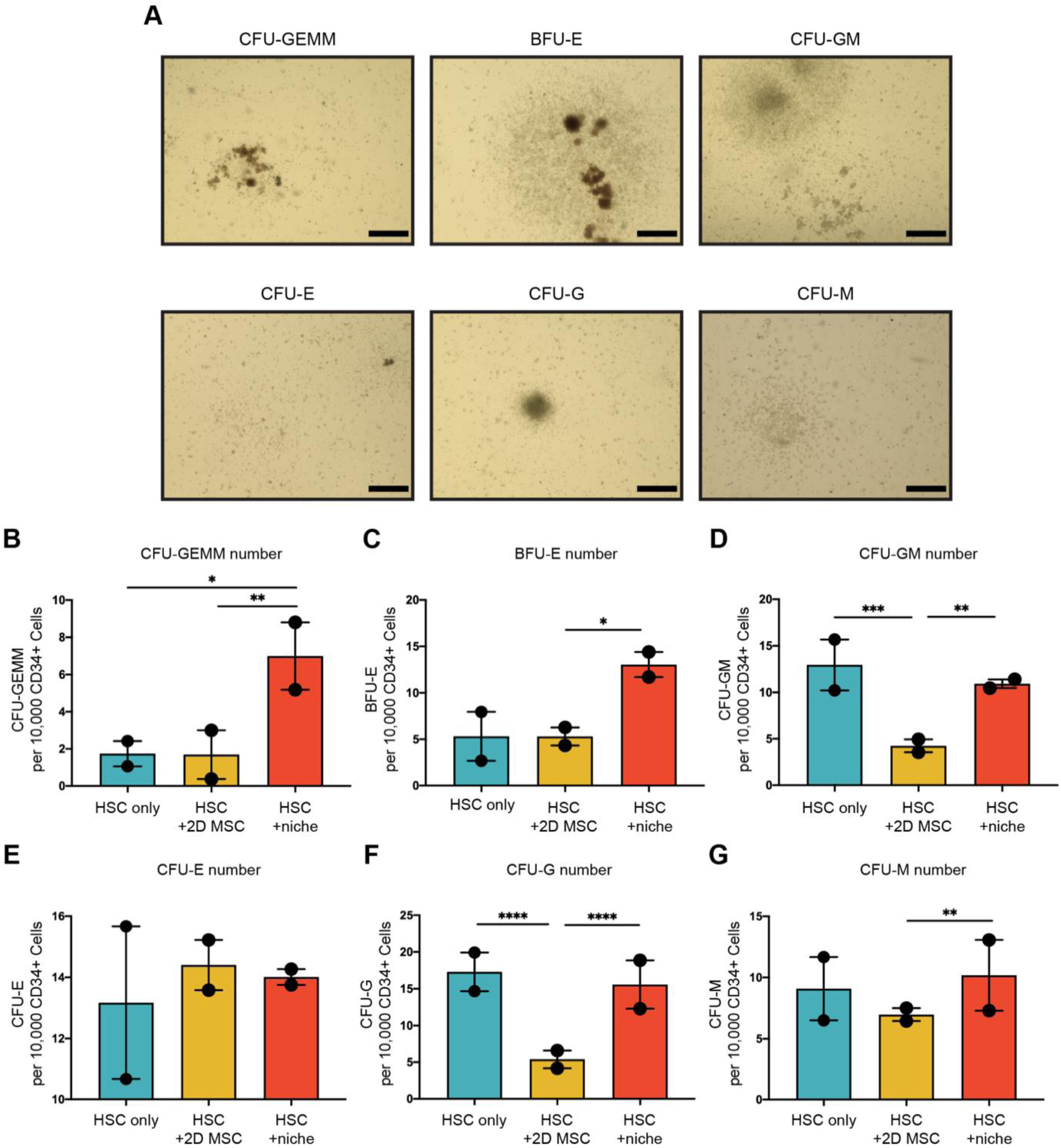
Representative images and CFU analysis. **(A)** Representative images of hematopoietic colony types in the CFU assay. CFU-GEMM, colony forming unit-granulocytes, erythrocytes, macrophages and megakaryocytes; BFU-E, burst forming unit-erythroid cells; CFU-GM, colony forming unit-granulocytes and macrophage; CFU-E, colony forming unit-erythroid cells; CFU-G, colony forming unit-granulocytes, CFU-M, colony forming unit-macrophages. These experiments were based on 2 independent donors with >3 technical repeats. Scale bar: 500 μm. Bar plots represent the colony numbers of **(B)** CFU-GEMM, **(C)** BFU-E, **(D)** CFU-GM, **(E)** CFU-E, **(F)** CFU-G and **(G)** CFU-M in each model. Mean ± s.d. based on n= 2 independent donors with >3 technical repeats. *, p<0.05, **, p<0.01, ***, p<0.001, ****, p<0.0001, ns, non-significant. P values were determined by ordinary two-way ANOVA with Tukey’s multiple comparisons test.

**Fig. S12.**
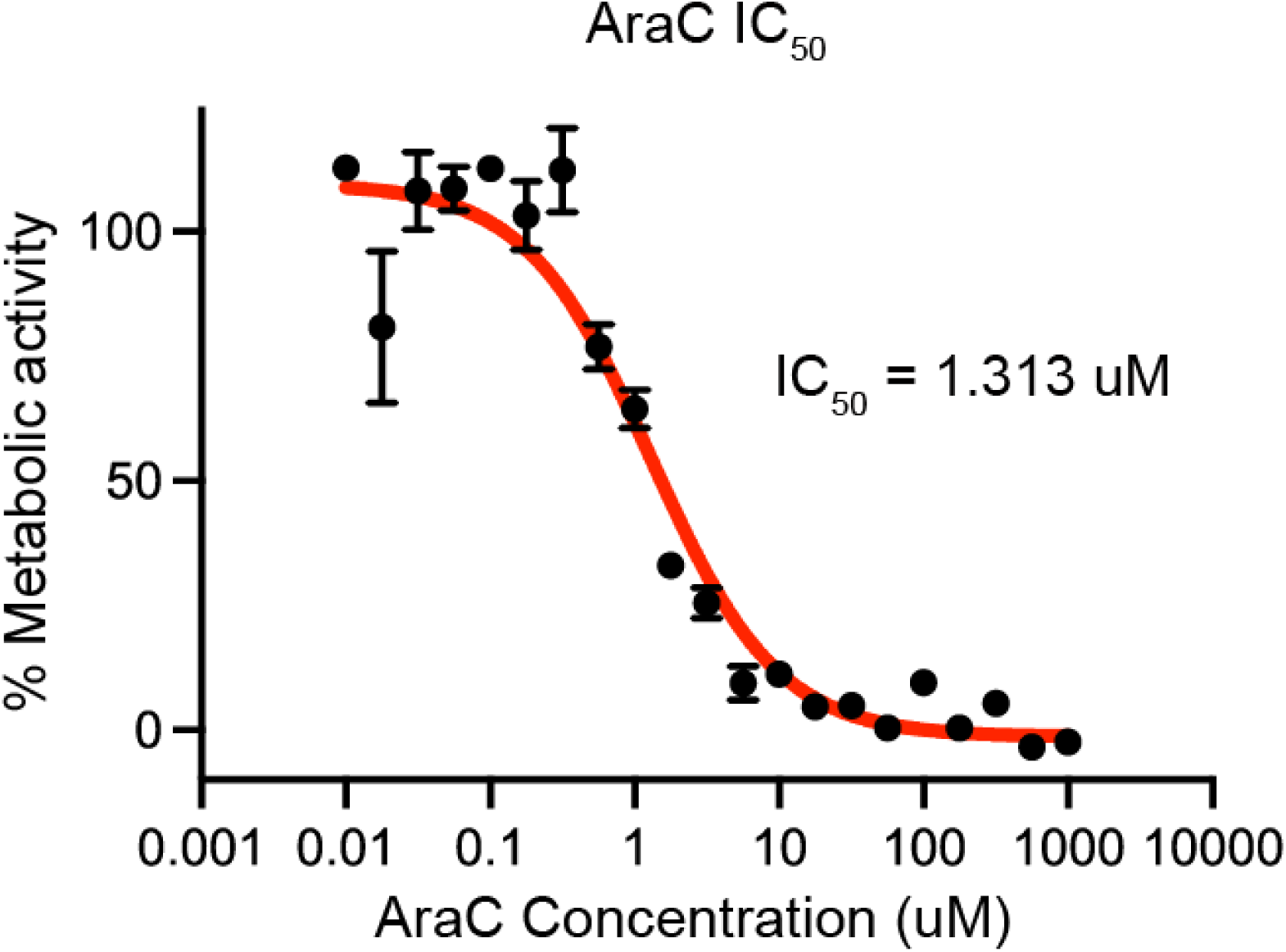
Dose-response curve of for THP1 exposed to AraC. The IC_50_ of THP1s to AraC at 72h using an Alamar blue assay. Mean ± s.d. based on n= 3 independent experiments.

**Fig. S13.**
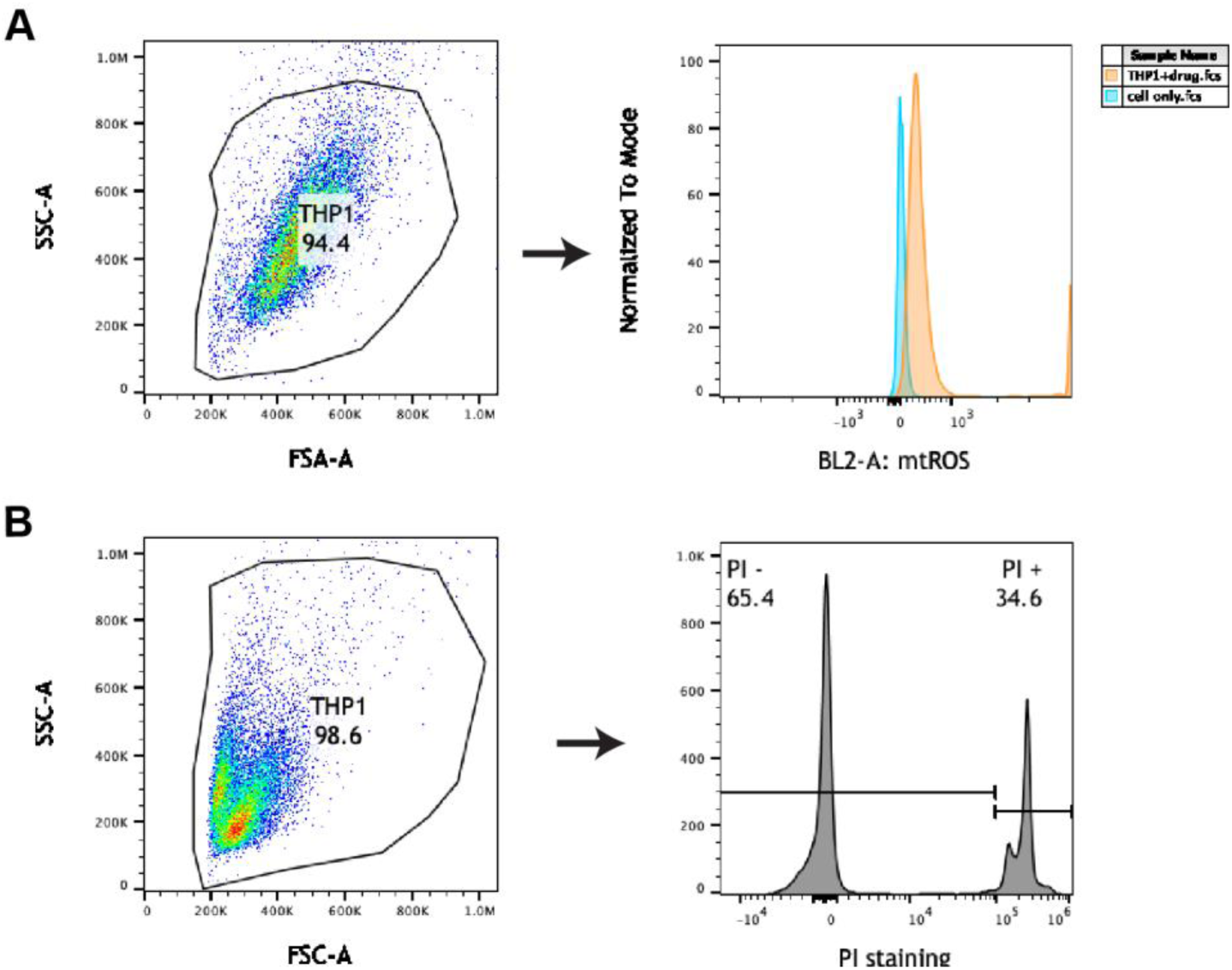
FACS gating strategy for mtROS and PI staining. **A,** FACS gating strategy for THP1 mtROS analysis. **B,** FACS gating strategy for THP1 PI staining.

**Fig. S14.**
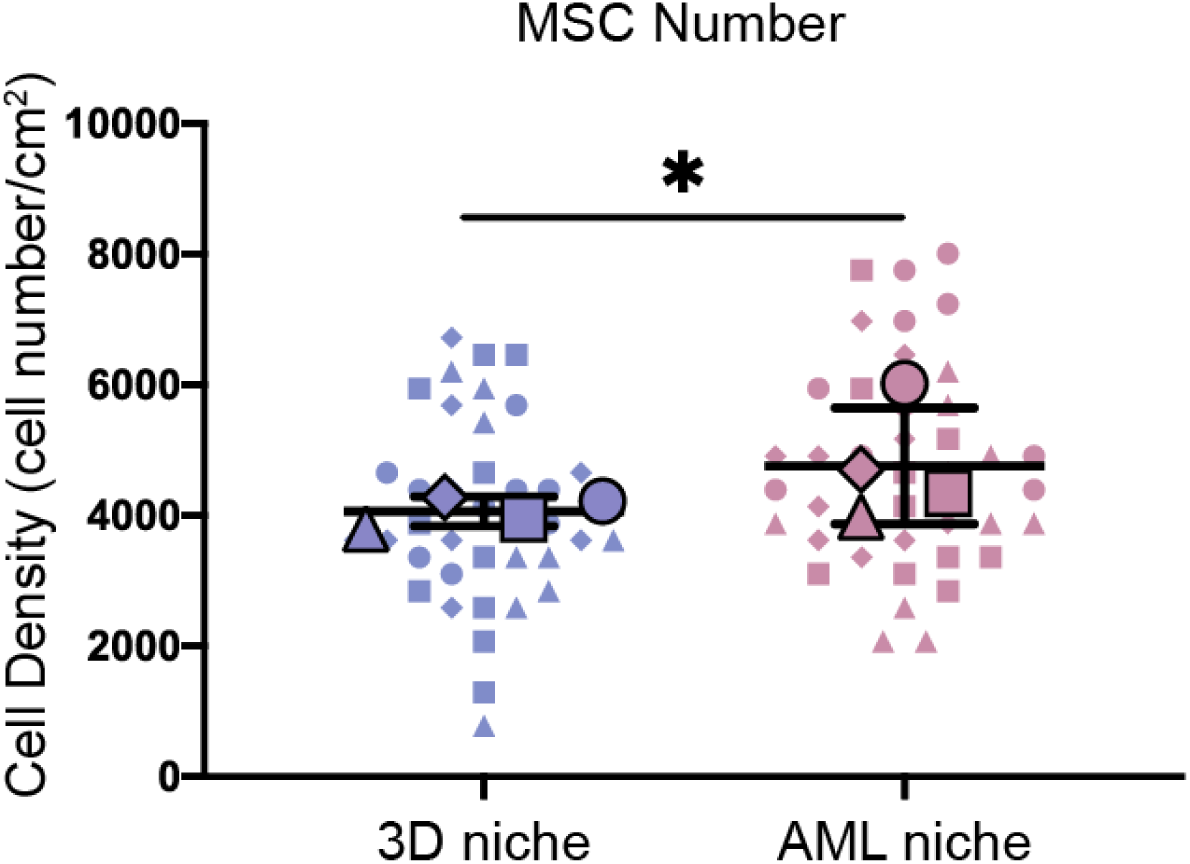
AML cells increased MSC proliferation. Live MSC counts (Hochest staining for live cell labelling) in the 3D niche model and AML niche model. Each shape represents each donor, each small shape represents each image field, and each large shape represents the mean from each donor. Mean ± s.d. based on n= 4 independent experiments with different donor cells. *, p<0.05. P values were determined by unpaired Student’s t-test was applied for two-group comparisons.

**Fig. S15.**
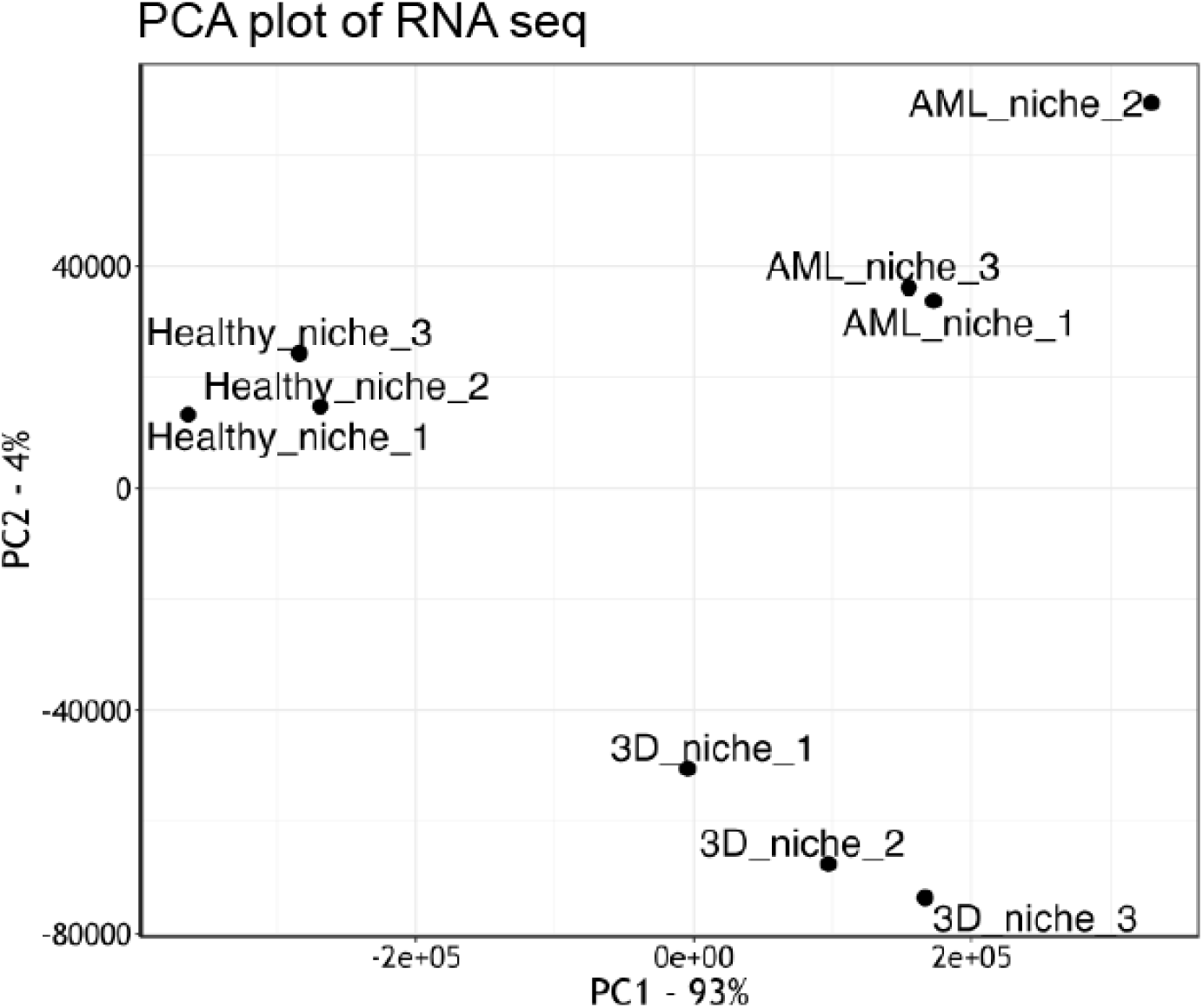
RNA seq of MSCs in the AML and healthy niches. PCA plot of transcriptional gene profile of MSCs from the 3D niche, healthy niche, and AML niche model.

**Fig. S16.**
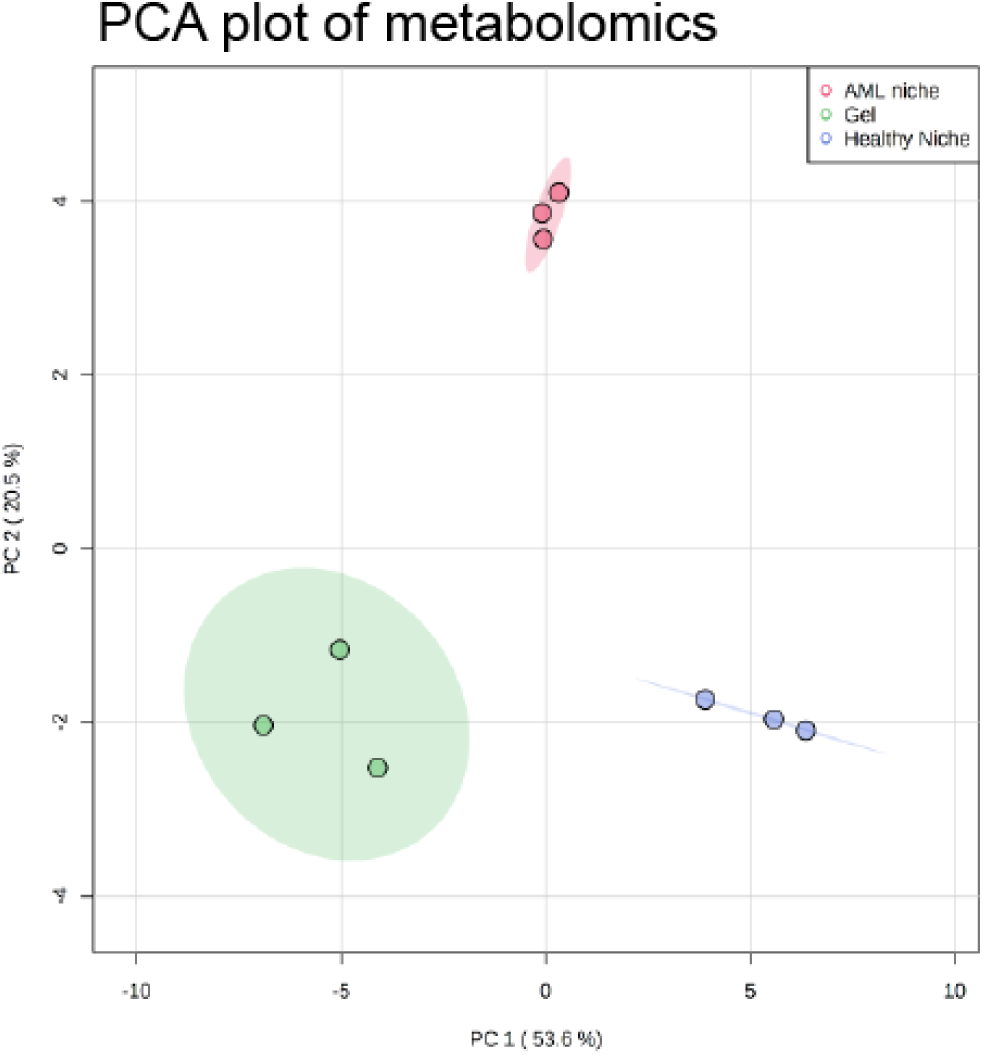
PCA plot of MSC metabolites from each niche model. PCA plot of the metabolites from MSCs from the 3D niche, healthy niche, and AML niche models. Experiments were based on n= 3 independent experiments with different donor cells.

**Fig. S17.**
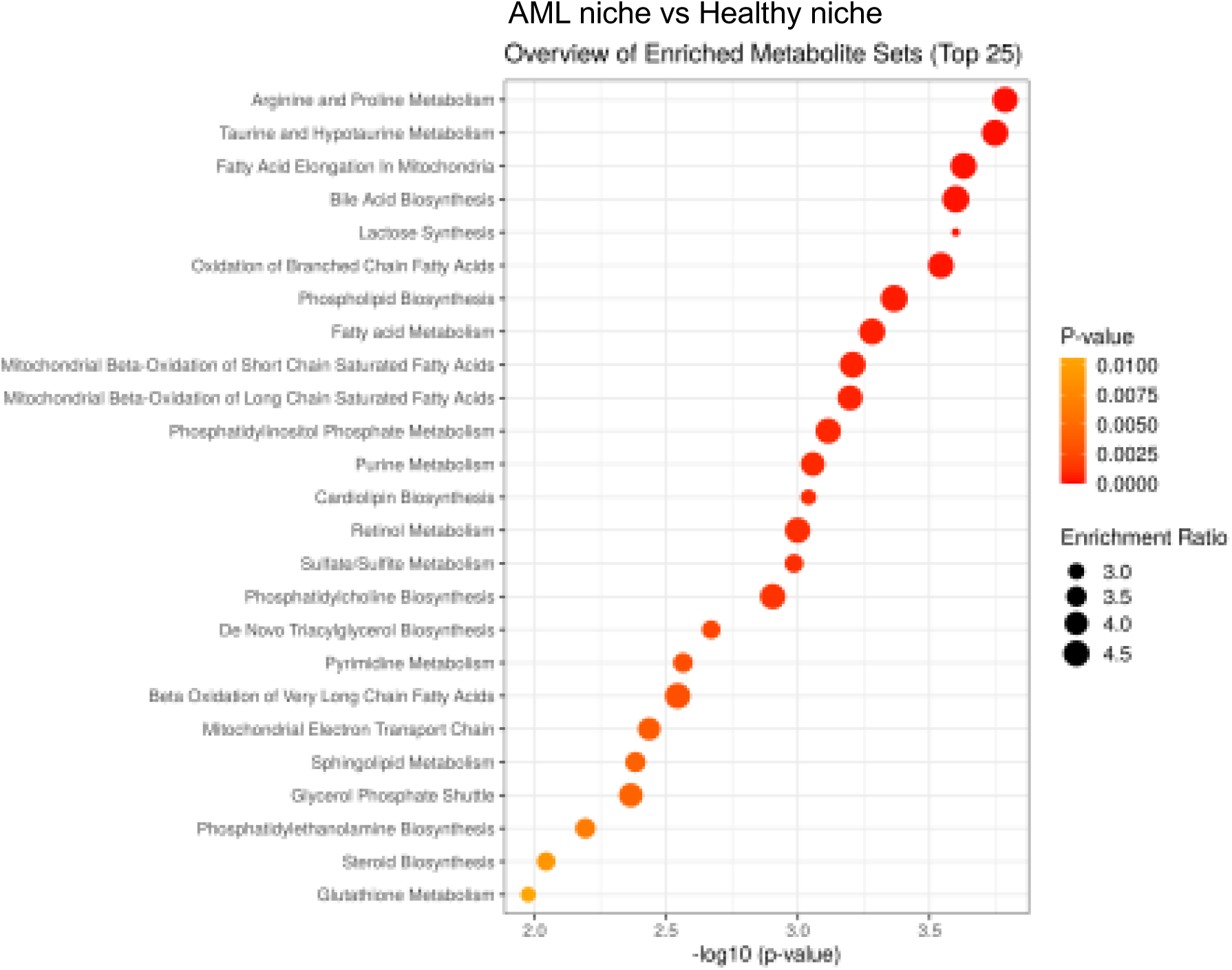
A larger version of Fig. 5K.

**Table S1.**
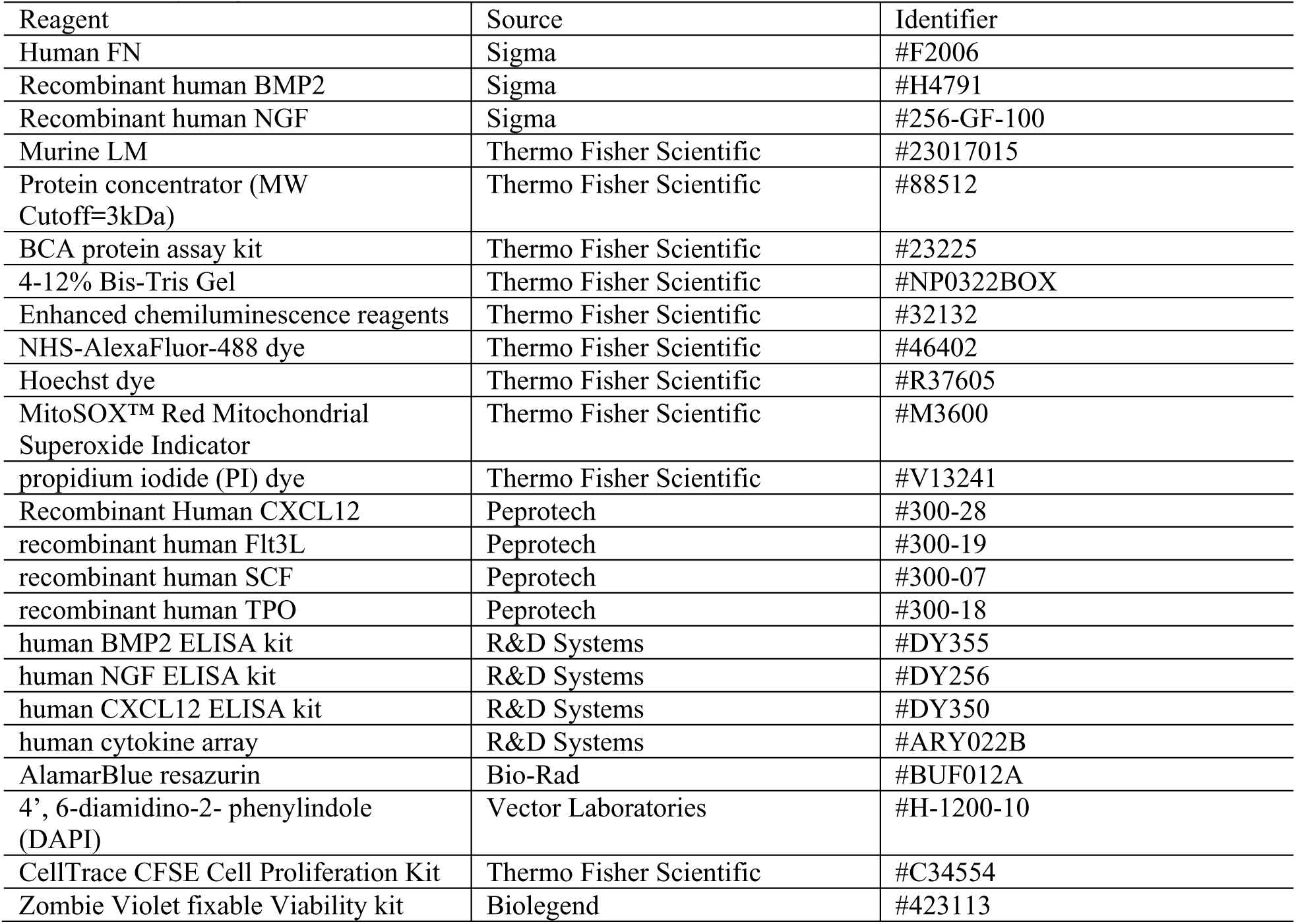
List of reagents used.

**Table S2.** List of antibodies used.

| Reagent | Source | Identifier | Dilution |
| --- | --- | --- | --- |
| Rabbit polyclonal anti-laminin antibody | Sigma | #L9393 | 1:200 |
| Mouse monoclonal anti-fibronectin antibody | Sigma | #F7387 | 1:200 |
| Mouse monoclonal anti-vinculin antibody | Sigma | #V9264 | 1:400 |
| monoclonal anti-CD45-FITC antibody | Thermo Fisher Scientific | #11-0459-42 | 1:50 |
| monoclonal anti-CD34-PE antibody | Thermo Fisher Scientific | #12-0349-42 | 1:100 |
| monoclonal anti-CD38-PE-Cy7 antibody | Thermo Fisher Scientific | #25-0388-42 | 1:200 |
| monoclonal anti-Lineage-FITC antibody | Thermo Fisher Scientific | #22-7778-72 | 5 ul/sample |
| monoclonal anti-CD34-PE antibody | Thermo Fisher Scientific | #12-0349-42 | 1:100 |
| AF-488 Phalloidin | Thermo Fisher Scientific | #A12379 | 1:200 |
| Rabbit monoclonal anti-phospho-FAK (Tyr397) antibody | Cell Signalling Technology | #8556S | 1:1000 |
| rabbit monoclonal anti-phospho-Myosin Light Chain 2 (Ser19) antibody | Cell Signalling Technology | #3671S | 1:1000 |
| mouse monoclonal anti-α-Tubulin (DM1A) antibody | Cell Signalling Technology | #3873 | 1:2000 |
| HRP-conjugated anti-mouse IgG antibody | Cell Signalling Technology | #7076 | 1:4000 |
| HRP-conjugated anti-rabbit IgG antibody | Cell Signalling Technology | #7074 | 1:4000 |
| Mouse monoclonal anti-P5F3 | Santa Cruz Biotechnology | #sc-18827 | 1:2000 |
| mouse monoclonal anti-OPN antibody | Santa Cruz Biotechnology | #sc-21742 | 1:50 |
| mouse monoclonal anti-YAP antibody | Santa Cruz Biotechnology | #sc-101199 | 1:100 |
| Rat monoclonal anti-HEP antibody | R&D Systems | #MAB4656 | 1:2000 |
| Mouse monoclonal anti-nestin antibody | Abcam | #ab22035 | 1:200 |
| rabbit monoclonal anti-VCAM antibody | Abcam | #ab134047 | 1:200 |
| Mouse anti-HFN7.1 antibody | Developmental Studies Hybridoma Bank |  |  |
| Gold particle-conjugated anti-rat IgG antibody | Aunion | #815.011 | 1:20 |
| IRDye 680RD secondary antibody | LI-COR |  | 1:800 |
| IRDye 800CW secondary antibody | LI-COR |  | 1:800 |
| CellTag 700 Stain antibody | LI-COR | #926-41090 | 1:500 |
| anti-mouse Texas red conjugated secondary antibody | Vector Laboratories | #TI-2000 | 1:50 |
| anti-rabbit Texas red conjugated secondary antibody | Vector Laboratories | # TI-1000 | 1:50 |
| rat IgG isotype antibody | Vector Laboratories | #TI-4000 | 1:20 |
| monoclonal anti-α5β1 antibody (JBS5) | Sigma | #MAB1969 | 1:20 |
| monoclonal anti-α6β1 antibody (GOH3) | Sigma | #MAB1378 | 1:20 |
| rat IgG isotype antibody | Vector Laboratories | #TI-4000 | 1:20 |

